# Loss of dominant caterpillar genera in a protected tropical forest

**DOI:** 10.1101/631028

**Authors:** Danielle M. Salcido, Matthew L. Forister, Humberto Garcia Lopez, Lee A. Dyer

## Abstract

Reports of biodiversity loss have increasingly focused on the abundance and diversity of insects, but it is still unclear if substantive insect diversity losses are occurring in intact low-latitude forests. We collected 22 years of plant-caterpillar-parasitoid data in a protected tropical forest and found reductions in diversity and density of these insects that appear to be partly driven by a changing climate and weather anomalies. The decline in parasitism represents a reduction in an important ecosystem service: enemy control of primary consumers. The consequences of these changes are in many cases irreversible and are likely to be mirrored in nearby forests; overall declines in the region will have negative consequences for surrounding agriculture. The decline of important tropical taxa and associated ecosystem function illuminates the consequences of numerous threats to global insect diversity and provides additional impetus for research on tropical diversity.

## INTRODUCTION

The impacts of global change are multi-faceted and ubiquitous *(1)* with major ecological and evolutionary consequences *(2)* that span aquatic and terrestrial ecosystems as well as a wide diversity of taxa and species interactions *(3)*. Much of global change research has focused on effects on single trophic levels, and despite an increased emphasis on interaction diversity in ecology *(4)*, relatively few studies have linked climatic variability to interaction diversity, ecosystem stability, and services of specific guilds, such as parasitoids. Past studies have also been geographically and taxonomically biased towards temperate ecosystems *(5–7)* and the subset of tropical studies are focused on vertebrates and focal tree species *(8)*. Despite the fact that 85% of global insect diversity resides in the tropics *(9)*, current analyses on their decline are primarily drawn from data from western, higher-latitude regions: United States, Great Britain and Europe *(10)*. Thus, although it has been clear for some time that a sixth mass extinction event is underway *(11)*, only recently have studies attempted to document declines in insect abundance and diversity in intact tropical forests by quantifying abundances of species within broad guilds *(12)*.

Documenting long term population trends and fluctuations in diversity in tropical insect communities is especially important because of an unjustified assumption that tropical communities are more stable *(13,14)* and therefore more resilient to multiple global change disruptions. Threats to insect diversity include climate change, habitat loss, fragmentation, invasive species, pesticides, and pollutants *(15–20)*, and the magnitude of these effects and associated levels of ecosystem resilience do indeed vary considerably across biogeographic regions. For example, changes in some climate parameters, such as mean annual temperature are most severe at the poles, and some of the most dramatic examples of biotic change have been observed at high latitudes, such as increased overwintering survival and voltinism in pest insects *(21, 22)*. In contrast, increases in extreme weather events will have complex and large effects on lowland tropical communities, where plant-insect food webs may be particularly sensitive because of highly-specialized trophic relationships relative to interactions at higher latitudes *(23)*. Furthermore, vulnerability of tropical communities to global change is exacerbated by the thermal constraints of tropical ectotherms *(24–26)* high degrees of endemism and high rates of tropical habitat loss *(27–30)*.

In general, reports on insect declines have mostly included cases where the causes are unspecified or unclear *(12, 31)*, or the consequences to ecosystem services have not been explored *(12, 16, 32)*. Here we contribute to understanding species declines and losses of biological interactions in a protected and well-studied tropical wet forest and examine potential losses of ecosystem function. The study area is La Selva Biological Research Station, Heredia Costa Rica (10° 26′ N, 83° 59′ W), a ~1600-ha patch of forest on the eastern Caribbean slope of the Cordillera Central, bordered by agriculture as well as the Braulio Carrillio National Park (Fig. 1A). We used data from 1997-2018 to examine changes in taxonomic diversity among larval Lepidoptera (“caterpillars”) and associated parasitic Hymenoptera and Diptera (“parasitoids”).

**Figure 1.**
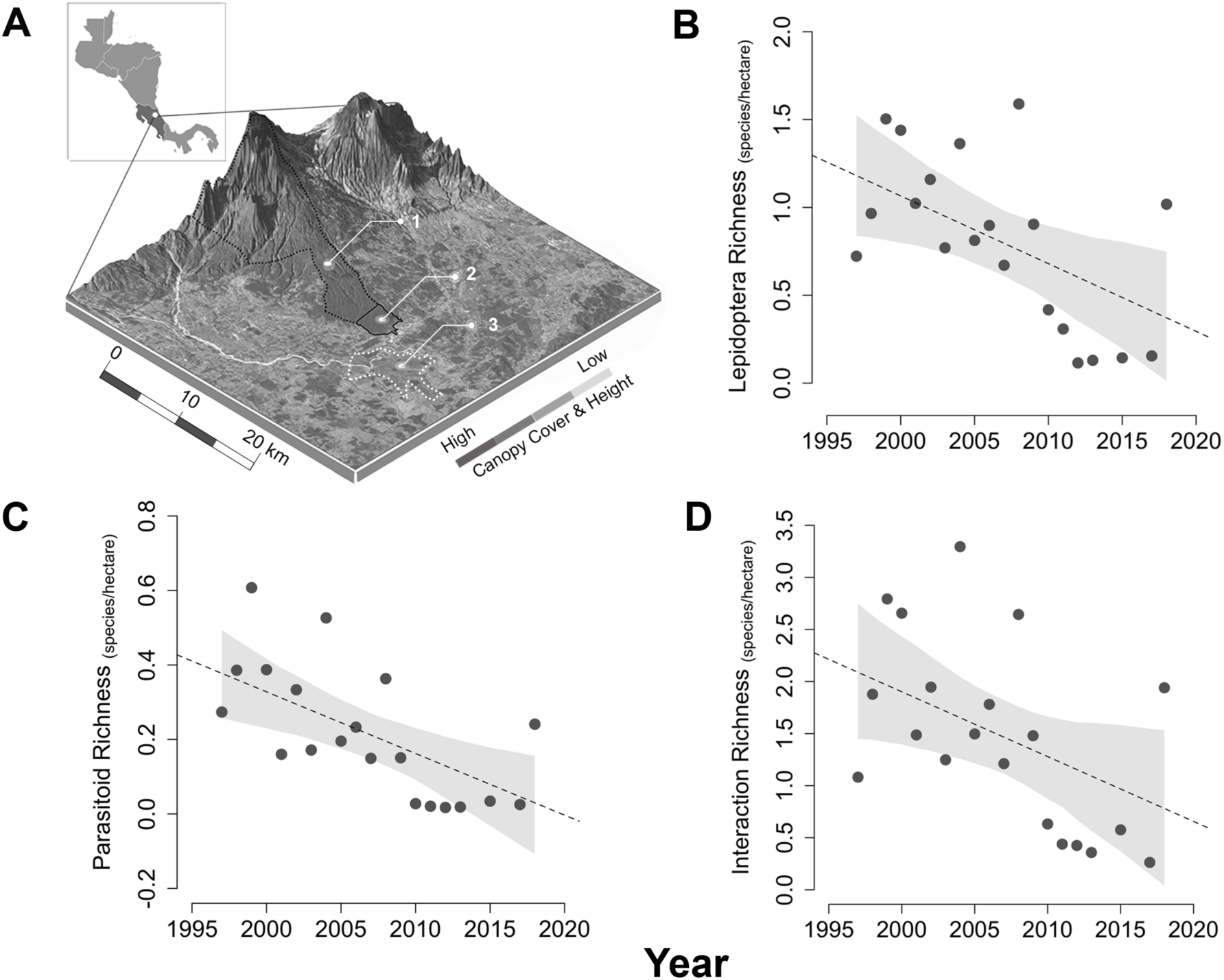
Insect declines across 22-years of sampling at La Selva Biological Research Station. Braulio Carillo National Forest (A.l) and surrounding areas, including La Selva (A.2), which has experienced declines in caterpillar diversity (β=-0.03, 95% credible intervals (CI) [−0.06,−0.01]) (B), associated parasitoid diversity (β =−0.02, [−0.03,−0.01]) (C) and interaction diversity (β =-0.07, [−0.13,−0.02]) (D) over the past 22 years (1997-2018). A large adjacent banana plantation is indicated by dashed white lines (A.3). Dotted lines on plots depicting declines are the best fit lines from Bayesian regression, with 95% credible intervals in gray.

## RESULTS

Our data reveal that declines in insect richness (Fig.1a,b) and diversity (Fig. S1–S3) are widespread across the two consumer trophic levels (caterpillar<u>s</u>: β=-0.03, 95% credible intervals (CI) [−0.06,−0.01], parasitoids: β =−0.02, [−0.03,−0.01]). These coefficients represent a 9.48% and 14.76% decline in species per hectare each year for caterpillars and parasitoids, respectively. Extrapolation of estimated declines to the full 1600 ha of La Selva yielded estimates for the number of species that have either been lost from the forest since the start of the study or have been reduced to sufficiently low density that they are no longer detected (which likely amounts to effective extirpation from the perspective of ecological interactions): we estimate 1056 fewer herbivore species (with 95% Bayesian credible intervals from 2112 to 352), and 704 fewer parasitoid species (from 1056 to 352). For the herbivores, for which we have the most data, we additionally used the first 5 years of data to estimate a baseline diversity (Chao estimator) from which the losses represent a 38.8% reduction (with credible intervals from 77.6% to 12.9%). In addition to declines in caterpillar diversity, frequencies of encounter for entire genera of caterpillars are decreasing: out of the 64 genera studied, 41% (26 genera) have an 80% probability of being in decline (i.e. at least 80% of the mass of the Bayesian posterior distributions were less than zero for the year coefficients in regressions for each of these genera) (Fig. 2, Table S1). Genera with greater than 95% probability of declines include: *Xylophanes (99%), Pantographa (97%), Emesis (96%), Dysodia (96%), Gonodonta (96%)* and *Hylesia* (95%).

**Figure 2.**
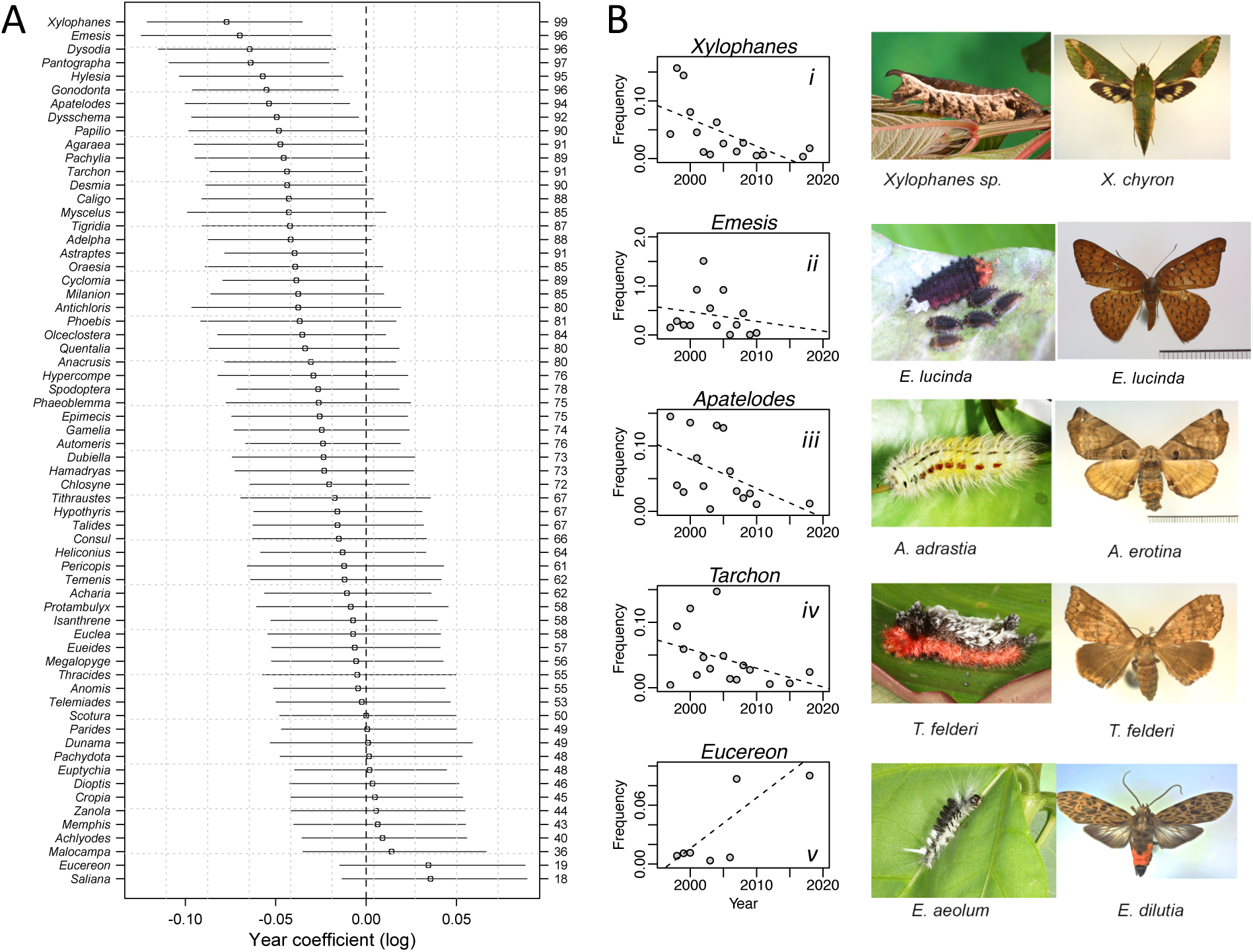
Genus-level patterns in caterpillar encounter frequencies across years. A) Point estimates for beta coefficients and associated 80% credible intervals (CI) for 64 genera that comprise a subset of all genera collected that met criteria for this analysis. Genus names are listed on the left margin and probabilities of a decline are on the right margin. Units of the year coefficient are the natural log of frequency per year. B) Frequency (untransformed) across years for select genera and representative larval and adult images.

Along with taxonomic declines, interaction diversity at La Selva is decreasing (β =−0.07, [−0.13,−0.02]; Fig. 1D): assemblages today have approximately 2,464 fewer unique interactions (30.9% reduction) than networks of interactions 22-years ago (Fig. 3A-B, Table S2, S3). Herbivore-enemy interactions were disproportionately affected, with over 77% of connections disappearing between herbivores and parasitoids when comparing networks of interactions in the first and last five years of the study. Losses in species and interaction diversity were paralleled by reductions in parasitism frequency, an important measure of natural biological control (β =−0.003, [−0.007,0.001], Fig.S5). Estimates for declines in parasitism are equivalent to −3% per decade which represents a ~6.6% decline during the study period. Further, the probability of a negative slope for overall parasitism across time was ~92%. Consistent with the hypotheses predicting the vulnerability of more specialized species to climate change *(33)*, declines in parasitism were greatest among hymenopteran parasitoids (−0.001 [−0.004,0.002]) compared to dipteran parasitoids (−0.00007 [−0.003, 0.003]). Overall this represents a −2.2% loss in parasitism by Hymenoptera compared to Diptera <0.2 % across the study period.

**Figure 3.**
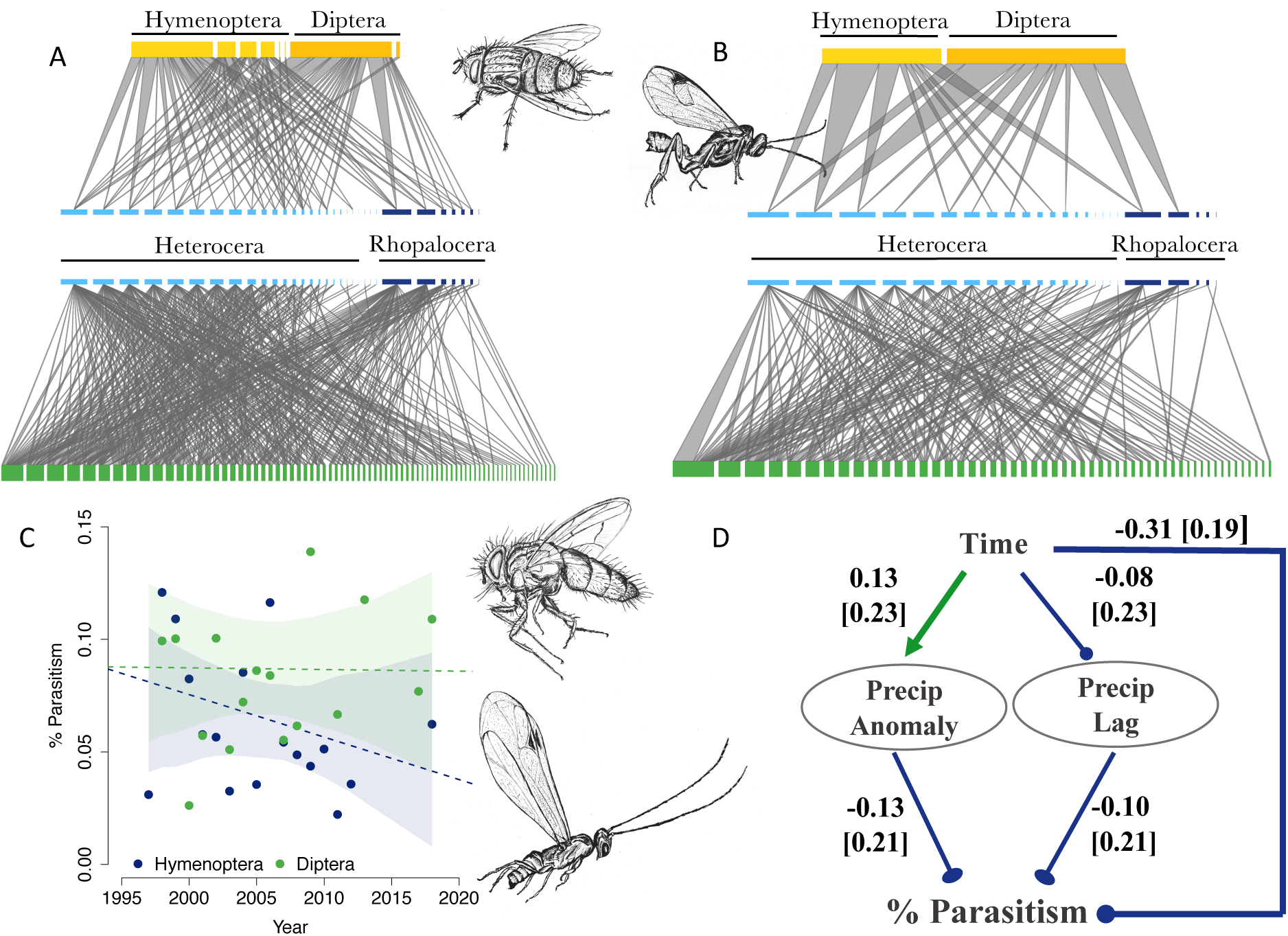
Patterns in plant-caterpillar-parasitoid interactions, climate, and parasitism across time. Tri-trophic networks illustrate host plants (green), caterpillars (blue), and associated parasitoids (yellow) for the first (A) and last 5 years (B) of the study. Nodes represent families within each trophic level and are grouped by suborder (Heterocera: light blue and Rhopalocera: dark blue) and order (Hymenoptera: light yellow and Diptera: mustard yellow) then ranked by node degree. Edge thickness represents relative link weights. Parasitism levels declined over time (C) (Hymenoptera: −0.001 [−0.004,0.002] and Diptera: −0.00007 [−0.003, 0.003] and these declines were associated with climatic changes (D). The structural equation model (good fit to the data: χ^2^ =0.887, p=0.346, df=1) illustrating these associations estimates effects of time on climate and time and climate on percent parasitism. Models for species richness and interaction diversity yielded similar coefficients as did models for parasitism with additional climate variables (Fig. S10–S11). Path coefficients are standardized and width of arrows are scaled based on magnitude of path coefficients. Standard errors are reported in brackets. Arrows represent positive associations and lines with circles represent negative associations. Parasitoid illustrations by M.L.F.

Environmental changes in and around the forest include persistent changes in land use and climate; annual temperature and precipitation are both increasing at La Selva (Fig. S6–S9, Tables S4, S5). In the last five years, precipitation anomalies have been larger than previous years and the number of positive temperature anomalies are increasing, with the greatest T_min_ and T_max_ anomalies on record occurring during the study period. Structural equation models (SEM) provided support for hypothesis that climate and climate anomalies have negative effects on species richness and ecosystem function. Precipitation anomalies and their one year time lag are among the most important factors causing lower parasitism frequency (χ^2^ =0.887, p=0.346, df=1; P > 0.05 indicates fit of the model to the data), following predictions of Stireman *et al.* 2005 *(33)* (Fig. 3D). Specifically, precipitation anomalies had a significant negative effect on percent parasitism (standardized path coefficient (hereafter, spc) = −0.13) as did the one year time lags of extreme precipitation events (spc=−0.1). Declines in richness are linked to changes in numerous climate variables. The best fit models provide support for the inference that declines in richness are caused by increases in precipitation and temperature anomalies (Fig. 4A, χ^2^ =0.044, p=0.833, df=3) and by increases in precipitation anomalies and maximum temperature (Fig. 4B, *χ*^2^ =0.273, p=0.67, df=2). Temperature anomalies had negative effects on all levels of richness (Fig. 4A, spc_caterpillar_=−0.17, spc_parasitoid_=−0.19, spc_interaction_=−0.18). Precipitation anomalies have more subtle negative <u>e</u>ffects on caterpillar and interaction richness and are associated with a similarly-slight increase in parasitoid richness (Fig. 4A, spc_caterpillar_=−0.08, spc_parasitoid_=0.06, spc_interaction_=−0.3). Rising maximum temperatures (Fig. 4B, spc=0.6) have caused relatively large decreases in all levels of richness (spc_caterpillar_=−0.24, spc_parasitoid_=−0.18, spc_interaction_=−0.19).

**Fig. 4.**
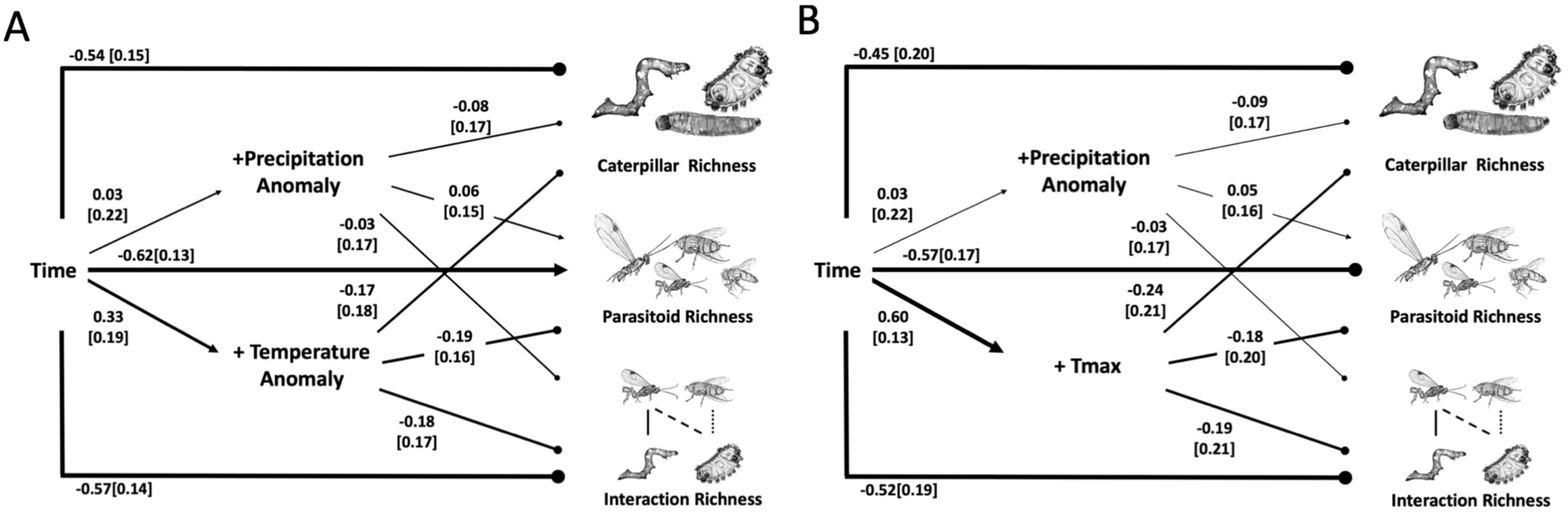
Structural equation models (SEM) estimating the effects of climate variables on caterpillar, parasitoid and interaction richness. A) Time (year) is an exogenous variable and the endogenous variables include richness, positive temperature anomalies and precipitation anomalies; model fit: χ^2^ =0.044, p=0.833, df=1. B) Similar variables as in the previous panel, but with average daily maximum temperatures instead of temperature; model fit: *χ*^2^ =0.173, p=0.67, df=1. Path coefficients are standardized and width of arrows are scaled based on magnitude of path coefficients. Arrows represent positive associations and lines with circle represent negative associations. Parasitoid illustrations by M.L.F. Caterpillar images by B.L.

## DISCUSSION

The dramatic declines reported here suggest that many caterpillars at La Selva will be losers and few will be winners in response to global change *(34,35)*, resulting in an overall reduction in the role of caterpillars as herbivores and as food for other animals. Compelling examples of winners and losers include the success of the genus *Eucereon*, which includes outbreak species *(36)*, and the failure of formerly common genera, such as *Emesis* (Fig. 2 & S4). The loss of common taxa are likely affecting the ecology of other organisms at La Selva -- declines in insectivorous vertebrate predators, including bats and birds within and near the forest, have already been attributed to reductions in arthropod prey *(37,38)*. Not only are these insects important prey, but the insects that are declining at La Selva are involved in numerous ecosystem processes, including parasitism, pollination, and plant consumption. In general, declines in caterpillars indicate an overall decline in environmental quality, as butterflies and other moth taxa are considered indicator taxa *(39)*.

As we documented at La Selva, one consequence of species extirpation or declines is the loss of interspecific interactions, which are the basis of ecosystem stability and ecosystem services *(40)*. Questions about loss of interaction diversity are largely absent from global change literature, due to a dearth of quantitative empirical data *(41–43)*. Reductions in species *(44)* and interaction diversity *(45)* can cause reduced ecosystem function via loss of functional redundancy, with likely cascading effects on natural biological control, pollination, plant diversity, primary productivity, and nutrient cycling. The declines in parasitism reported here can be extrapolated to an impressive 30% drop in parasitism over the next 100 years, which is a major loss of a key ecosystem service that prevents damaging outbreaks of herbivorous insects *(33)*. Losses of species and trophic interactions of this magnitude are particularly relevant in areas with intensified agriculture, where the global economic contribution of biological control is now estimated at $1.56 trillion per year *(46,47)*. The tropics currently have the greatest rates of agricultural expansion and tropical agriculture is expected to expand by at least 50% by 2050 *(48, 49)*. For the continually expanding agricultural areas surrounding La Selva, parasitoids are essential for biological control; for example parasitoids are an effective control of herbivores in banana *(50)*, and over 10,000 ha of land surrounding La Selva are banana plantations with one of the largest plantations situated <3km from La Selva (Fig. 1A).

Climate-driven declines in parasitism were paralleled by climate-driven declines in insect and interaction richness. Anomalies in precipitation (extreme wet events) and temperature (warmer than average episodes) are occurring at an increasing rate which is consistent with the idea that tropical ecosystems are facing increasing climatic variability and extremes *(51, 52)*. Extreme climate weather events such as flooding (positive precipitation anomalies) and drought (positive temperature anomalies) negatively affected insect richness and associated interactions in our study area. In addition to an elevated frequency of weather anomalies, maximum daily temperatures are increasing and appear to have a negative effect on the observed richness of caterpillars, parasitoids, and interactions. Although it is unclear if tropical endotherms will be more sensitive to changes in temperature relative to higher-latitude species *(53)*, our results confirm that insect food webs are affected by warming conditions. Increases in minimum temperature (Fig. S11) are also contributing to declines in insect richness which is consistent with responses reported in temperate butterfly systems *(54)*. In all models, time had the second strongest (direct negative) effect on richness and parasitism frequency compared to other predictors, suggesting that other unmeasured global change variables also contribute to insect declines at La Selva and further research is needed in this area.

Declines in populations of plants and animals, extinctions, and loss of ecosystem function are defining features of the Anthropocene *(11)*. From a general Bayesian perspective in which new results are used to update prior knowledge *(55)*, additional corroborations of these Anthropocene-associated losses are useful in that they provide more precise estimates of decline probability for specific taxa, regions and ecosystems. Although insect declines have been the subject of recent high profile studies *(8,31)*, the taxonomic and geographic breadth of the phenomenon is not without controversy *(32)* and reports have been rare from the planet’s most species-rich ecosystems. Thus we suggest that the results reported here strengthen the growing probability that insects are facing what indeed may be a global crisis. The hard work that still faces ecologists is to try to figure out which traits and habitats most expose species to risk, while the challenges for taxonomists and natural historians are to discover and describe new species and interactions before they disappear. All scientists should be considering how to use existing data to focus on the most imperiled taxa, ecosystems, and biogeographic regions. Tropical wet forests are clearly one biome requiring more precise estimates of species declines and a better understanding of determinants of these declines. For La Selva, our results are consistent with the hypothesis that climate change in lowland tropical forests is causing declines in species and entire genera of herbivores as well specialized parasitoids. Although such multi-trophic connections are not frequently studied in the context of global change, if results such as ours are widespread, then cascading results to other guilds and trophic levels can be expected *(56)* and warrant immediate concern and management effort.

## METHODS

### Study sites and sample methods

We collected plant-caterpillar-parasitoid interaction data within La Selva across all seasons. Seasonality is marked by a wet season from May to December and a brief dry season from January to April; peak rainfall occurs in June-July, and March is the peak dry month. Samples were collected as a larger rearing program cataloguing plant-herbivore-parasitoid associations across the Americas *(57, 58)* from 1995 to present. We limited our results to records starting in 1997 up to 2018, and we excluded 2014 and 2016 because sampling days did not meet our minimum criteria of 20 sampling days/year. We sampled externally feeding (including shelter builders) caterpillars from their host plant and reared them to adult moths or parasitoids. Caterpillars were located opportunistically by visual inspection along trail transects (distance varies between 50-3000m and select transects were continuously sampled across years), or exhaustively sampled in 10m diameter plots (149 plots total) by staff scientists, graduate students, parataxonomists, and teams of Earthwatch volunteers and students. Due to variable sampling days across years, we weighted observed values by sampling effort. Sampling effort was calculated as the number of volunteer and staff days of sampling multiplied by the average area in square meters covered by each person in a 10-day sampling period (4000 m^2^); hence, observed diversity and frequency is presented and analyzed in models as species equivalents or frequencies per hectare per year. We excluded *Eois* (Geometridae) and *Quadrus* (Hesperiidae) from all analyses because these focal genera present a bias in the rearing dataset due to focused collection for ancillary studies.

## Rearing Methods & Processing Data

For our ongoing interaction diversity survey, collected larvae are given a unique voucher code that associate them with their host plant species. Caterpillars are reared individually in plastic containers or bags with a sample of hostplant. Species identifications are made initially by parataxonomists to lowest taxonomic level or morphospecies and verified by taxonomic experts or by referencing voucher specimens and image libraries. Some morphospecies are confirmed using mtDNA COI sequences, others by examining a mix of morphological characters, and others using genomic data. For the remaining species without morphospecies designations we assign morphotypes based on feeding relationships – morphologically distinct caterpillars from the same family utilizing the same host family are designated a unique morphotype. This method is likely a conservative means to assigning species names, especially for tropical species *(23, 58)*. Voucher specimens are sent to collaborating institutions including universities and museums (see www.caterpillars.org for a list of participating institutions)

## Patterns in Diversity, Parasitism, Climate Variables

### Abundance & Diversity

We quantified species abundances and unique interaction frequencies for each year to compare patterns in diversity and abundance across time. We also aggregated the data to examine annual genus-level frequencies of caterpillars to evaluate declines of higher taxa; genus-level abundance data analyses only included genera with ≥5 years of data and sampling that extended to 2010. Results are reported for the 64 caterpillar genera that met these criteria. To obtain values of interaction diversity, we modified a community matrix such that rows were comprised of years and columns the unique interactions. Interactions were comprised of bi-trophic (plant-herbivore) and tri-trophic (plant-herbivore-enemy) interactions and each matrix cell represented annual frequencies of those interactions.

Diversity was calculated using Hill numbers, and values were interpreted as interaction or species equivalents *(59,60)*. Hill numbers vary as a function of the parameter q and indicate the sensitivity of the index to rare species, and q=0, q=1 and q=2 are equivalent to species richness, Shannon’s diversity, and Simpsons diversity, respectively. We used functions provided in Chao *et al.* (2015) *(60)* to calculate Hill numbers. Results for q=0 are reported in the main text and q=1,2 in the supplemental information. To obtain estimated percent herbivore loss we quantified mean (Chao estimated) diversity for the first 5 years of data and subtracted species decline estimated from beta coefficients of the linear models of diversity across years.

### Climate Variables

Climate variables were calculated as annual means of daily precipitation and average, minimum and maximum temperatures. We used meteorological data acquired from weather stations within La Selva from 1983-2018 *(61)*. Temperature is reported as degrees Celsius (°C) and precipitation in millimeters (mm). To examine effects of extreme weather events and climate variability on patterns of diversity, we calculated anomalies and the coefficients of variation (CV) for each precipitation and temperature variable. Precipitation anomalies were calculated as the sum of daily values exceeding 2.5 standard deviations (sd) of the annual mean. Similarly, for temperature anomalies we used 2sd. The coefficient of variation was calculated as the ratio of standard deviation to the annual mean. We used simple linear regression to evaluate patterns among each climate variable across time and with respect to each season in the supplemental figures.

### Evaluating Patterns in Network Structural Properties

We pooled interaction data to the family level for the first (1997-2001) and last (2012-2018) five years of collection to illustrate changes in tri-trophic network structure. For each network we calculated node degrees and relative edge weights and reported link and node richness for each trophic level (Table S2).

### Parasitism Frequency

Percent parasitism was calculated as the ratio of parasitism events to all successfully emerged adults (caterpillars plus parasitoids) for each month from 1997-2018. We examined monthly trends across time to account for intra-annual and seasonal variation in tropical population dynamics. Excluded from analyses were months with zero parasitism or zero eclosed caterpillars as well as months without a number of adults that exceeded the 1^st^ quantile (Q_1_) of the distribution of total adults (IQR=12-103).

## Statistical Models

### Bayesian Models

We used Bayesian linear models to estimate coefficients for change over time for diversity of caterpillars, parasitoids, and interactions, as well as parasitism frequency. Models were fit for total parasitism and separately for specialized (Hymenoptera) and non-specialized (Diptera) orders of parasitoids. Models were fit in JAGS (version 3.2.0) utilizing the rjags package in R *(62)* using (for each analysis) two Markov chains and 1,000,000 steps each; performance was assessed through examination of chain histories (burnin was not required), effective sample sizes and the Gelman and Rubin convergence diagnostic *(63)*. Response variables were modeled as normal distributions with means dependent on an intercept plus predictor variables (either year alone, or year plus climatic variables), and vague or minimally-influential priors as follows: priors on beta coefficients (for year and climatic variables) were normal distributions with mean of zero and precision of 0.01 (variance = 100); priors on precisions were modeled as gamma distributions with rate = 0.1 and shape = 0.1. All data was z transformed prior to analysis. An additional hierarchical model (with vague priors as already described) was used to estimate change across years in the frequency of observations of individual caterpillar genera, with the year coefficients (and intercepts) estimated for each genus separately (as the lower level in the hierarchy) and simultaneously across all genera (the response variable for this analysis was log estimates from posterior distributions for beta coefficients, as well as 95% credible intervals for the diversity models and 80% intervals for the hierarchical model. We used the more liberal calculation of intervals for the latter in the interest of minimizing type II error in a situation involving the decline of entire genera (i.e., we would rather risk the possibility of erroneously inferring decline as opposed to mistakenly concluding that a declining taxon is stable). As a complementary measure of confidence not dependent on an arbitrary cutoff for importance, we calculated (for the beta coefficients estimated for each genus) the fraction of the posterior distribution less than zero, which can be interpreted as the probability that a genus has been observed with decreasing frequency over time.

### Structural Equation Models

We used Structural Equation Modeling (SEM) to test causal hypotheses that evaluated the effect of climate and time on taxonomic and interaction richness and parasitism. We used the global estimation method in the lavaan package v.0.6-3 *(64)* in R v 3.5.3 to generate 3 models. Models evaluated casual relationships among caterpillar richness, parasitoid richness, interaction richness, parasitism levels, and climate variables. Climate variables included: Tmin and Tmax and their anomalies and precipitation anomalies. A priori expectations of relationships between climate variables and parasitism *(33)* guided tests of causal hypotheses that parasitism levels are determined by precipitation and its one-year lag. Model fit was assessed using χ2 values and models were compared using Akaike information criteria (AIC). We reported standardized path coefficients and illustrated the SEM results in a path diagram.

## ACKOWLEDMENTS

We thank Earthwatch Institute and volunteers, UNR Plant-Insect Group, G. Gentry, J.O. Stireman, S. Shaw, J. Whitfield, J. Miller, J. Brown, L. Richards, A. Smilanich, J. Elliot, R. Parry, T. Davis, Z. Bousum, B. Carranza, D. Brenes & O. Vargas for substantive contributions to this work.

## Author Contributions

D.M.S., L.A.D., H.G.L. conceived project and supervised field work. D.M.S., L.A.D., M.F. analyzed data and wrote manuscript. Parasitoid illustrations by M.L.F. and caterpillar illustrations by B. Lopez.

## Data and materials availability

All data needed to evaluate the conclusions in the paper are available upon request.

## SUPPORTING INFORMATION

**Fig. S1.**
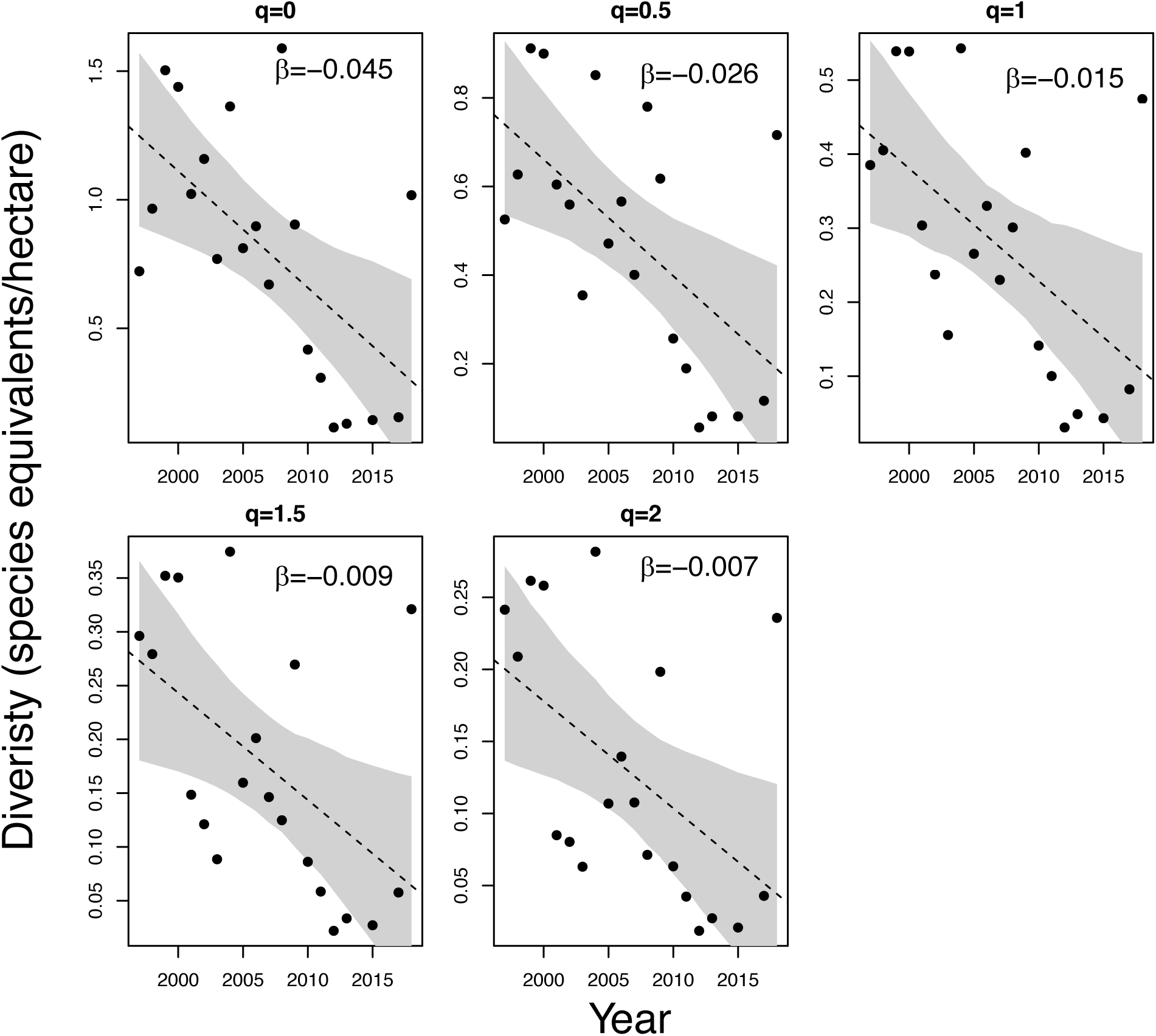
Lepidoptera diversity measured as effective number of species/hectare across sampling years. Each panel represents Hill numbers between 0 and 2. Beta coefficients were estimated using Bayesian linear models and shaded areas represent 95% credible intervals.

**Fig. S2.**
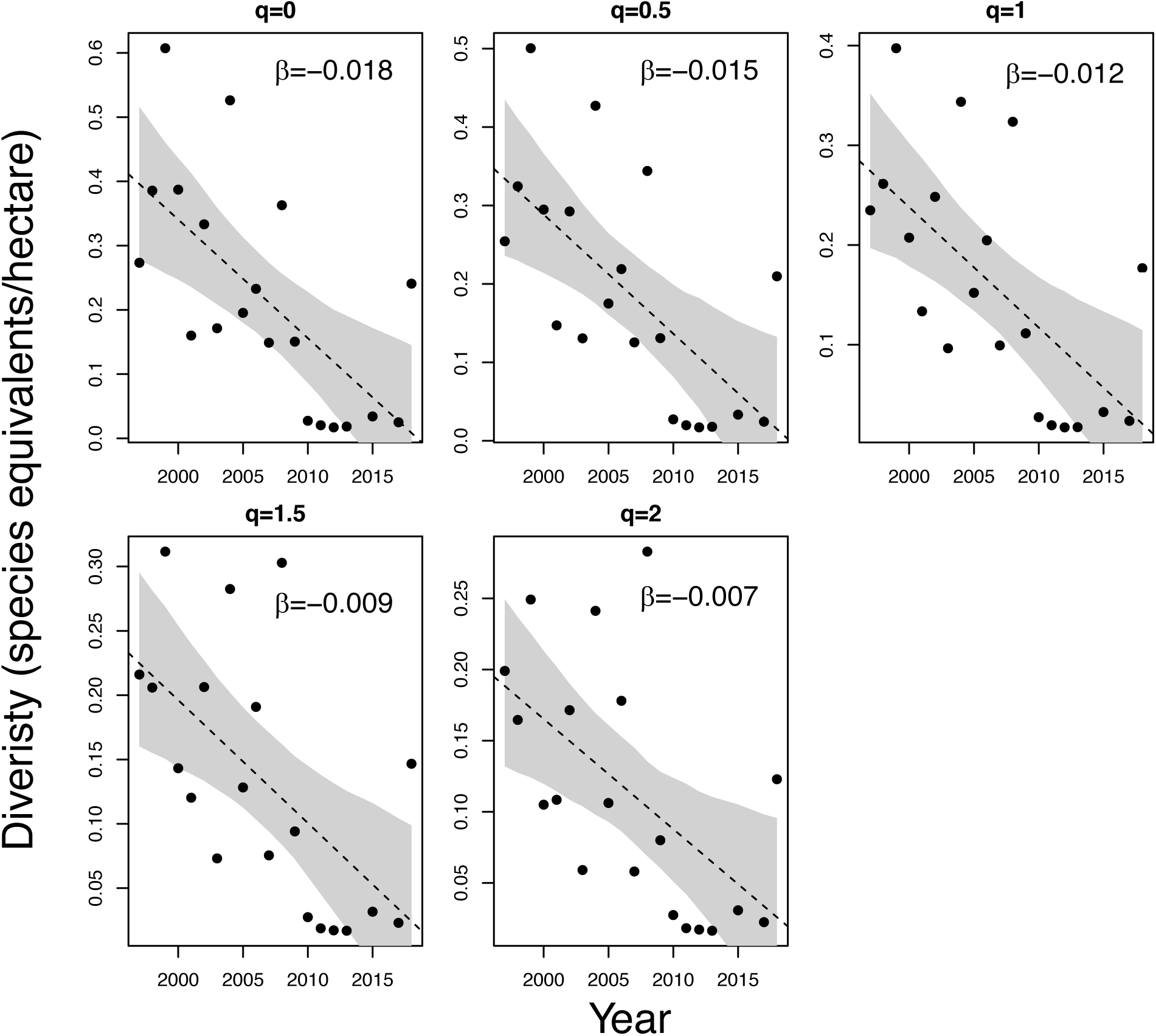
Parasitoid diversity measured as effective number of species/hectare across sampling years. Each panel represents Hill numbers between 0 and 2. Beta coefficients were estimated using Bayesian linear models and shaded areas represent 95% credible intervals.

**Fig. S3.**
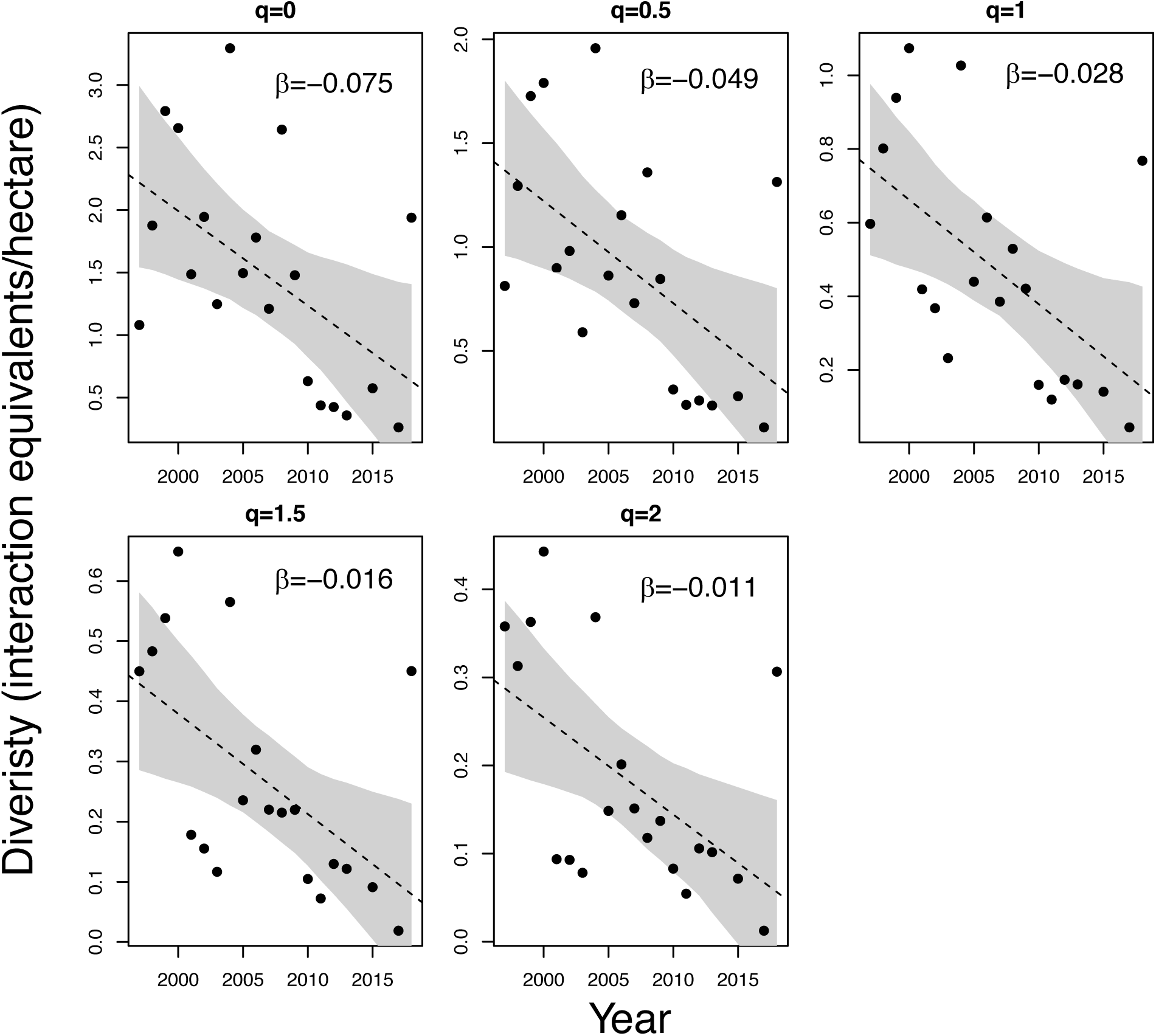
Tri-trophic interaction diversity measured as effective number of interactions/hectare across sampling years. Each panel represents Hill numbers between 0 and 2. Beta coefficients were estimated using Bayesian linear models and shaded areas represent 95% credible intervals.

**Fig. S4.**
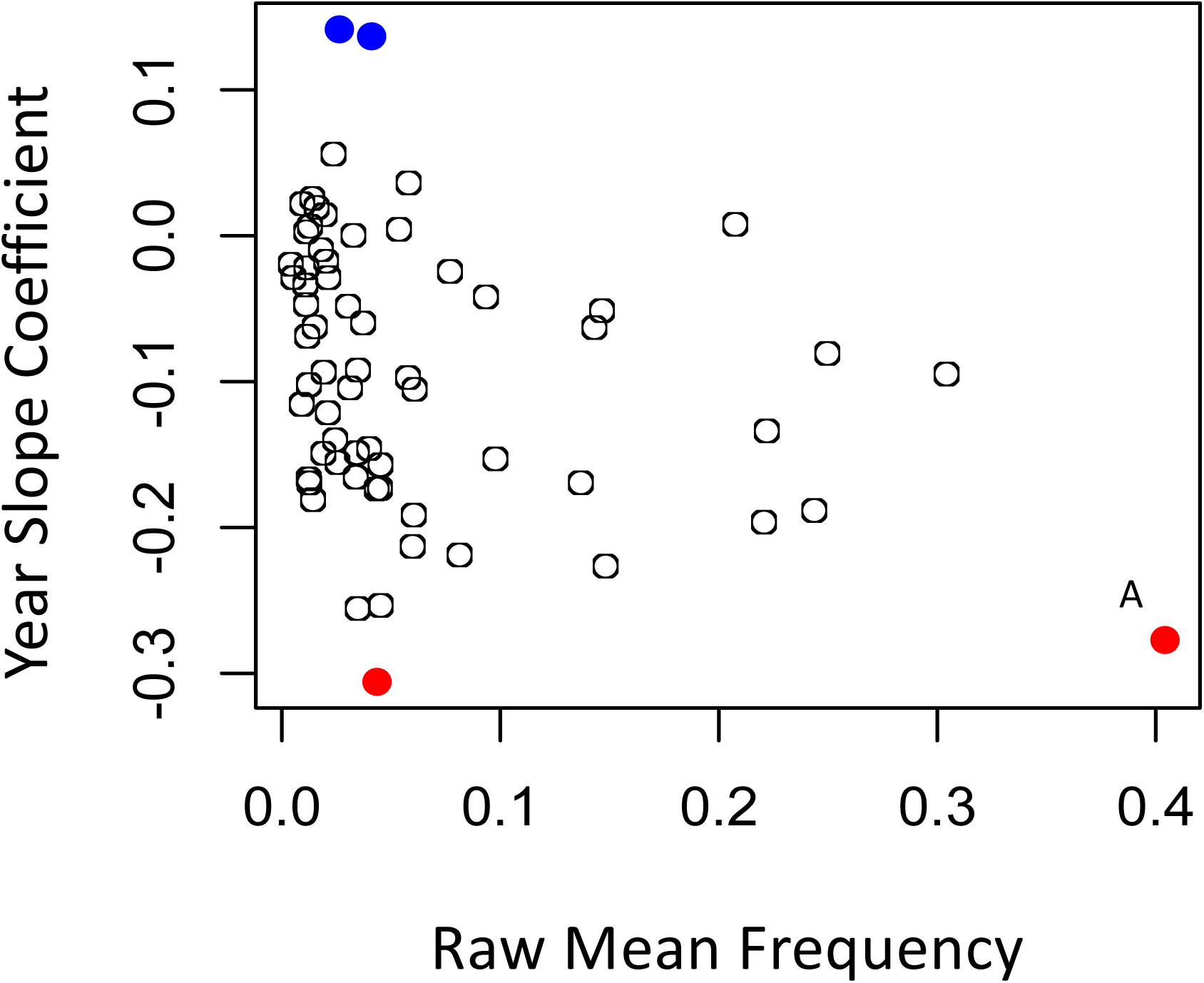
Beta coefficients for years plotted against raw frequencies of observation for 64 Lepidoptera genera. The blue points the two biggest ‘winners’ (*Euceron* and *Saliana*) and the red points are our two biggest ‘losers’ (*Emesis* (A) and *Xylophanes*).

**Fig. S5.**
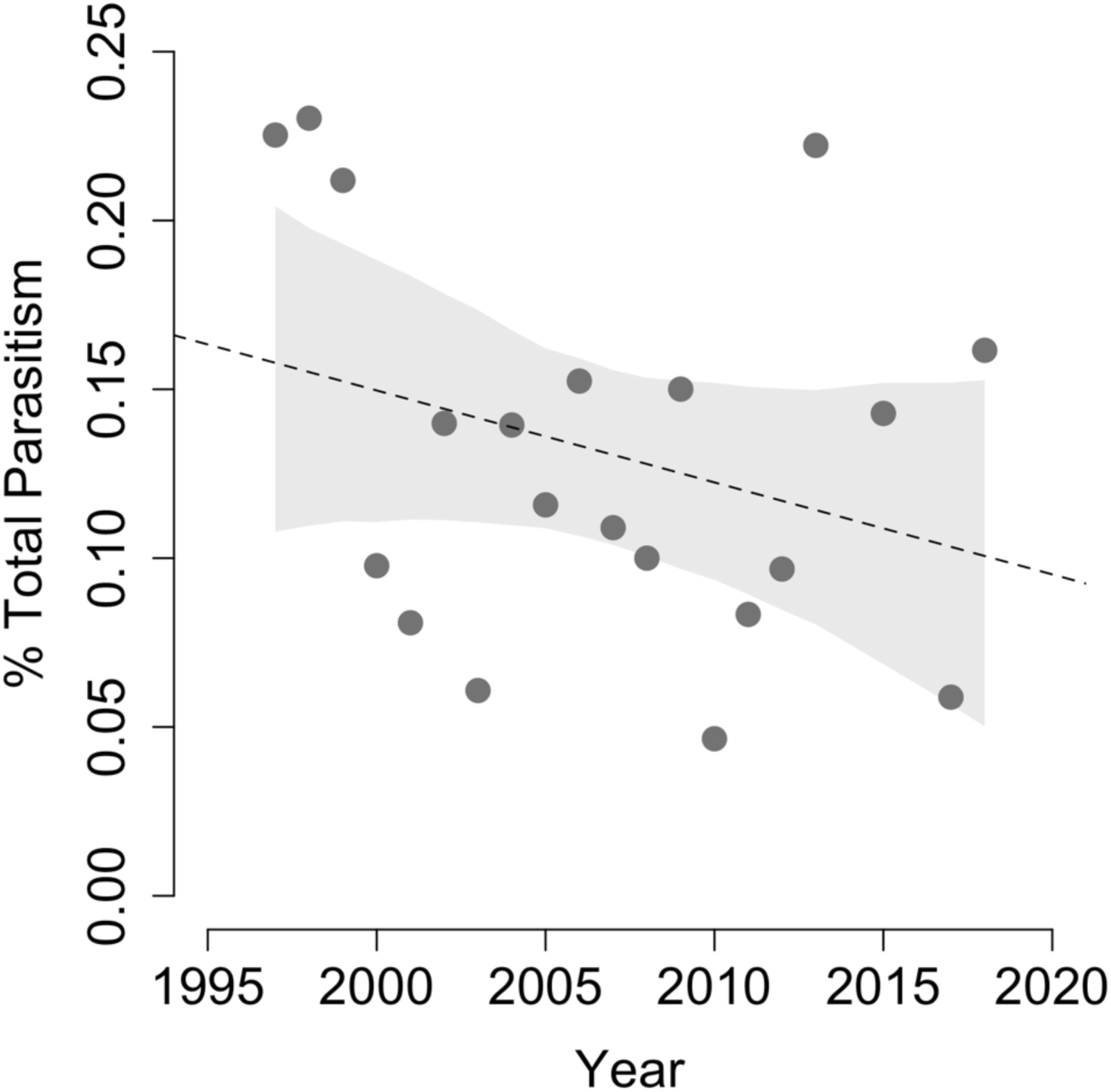
Parasitism frequency at La Selva across years of study (1997-2018). A Bayesian linear model was used to estimate beta coefficient (β =−0.003, [−0.007,0.001]); the shaded area displays 95% credible intervals.

**Fig. S6.**
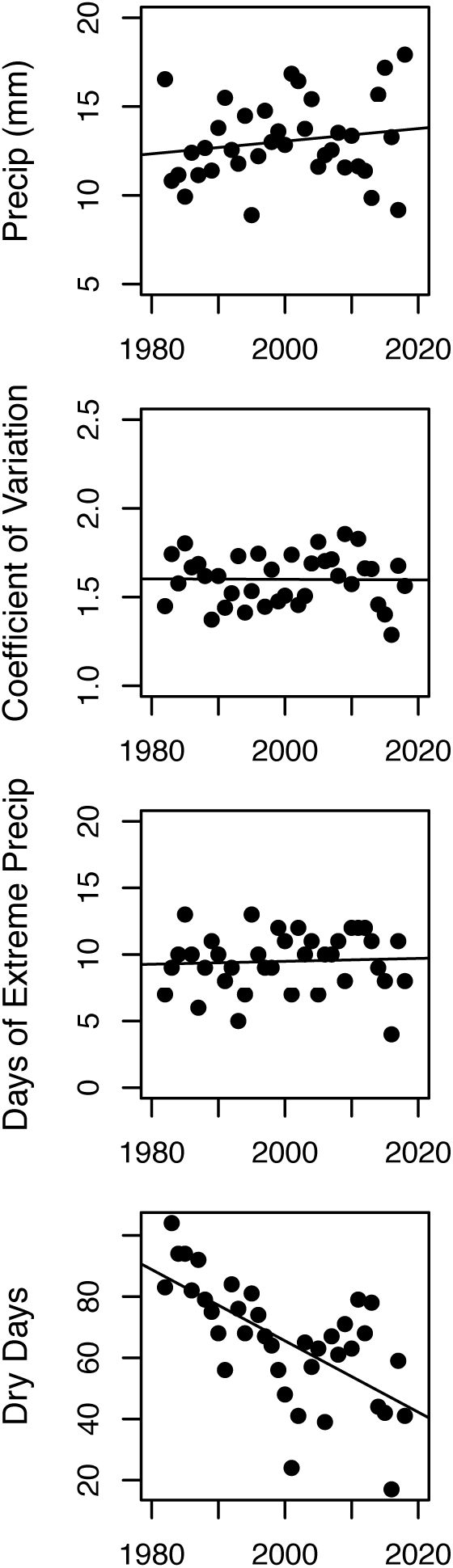
Patterns in precipitation variables at La Selva from 1982-2018. Each point represents a year of data. Precip (mm) is c the annual mean of daily precipitation<u>, t</u>he Coefficient of Variation (CV) in precipitation is intra-annual CV<u>, d</u>ays of extreme precipitation are counts of daily precipitation exceeding 2.5 SD of annual mean precipitation, and dry days were calculated as total days within a year with zero rainfall.

**Fig. S7.**
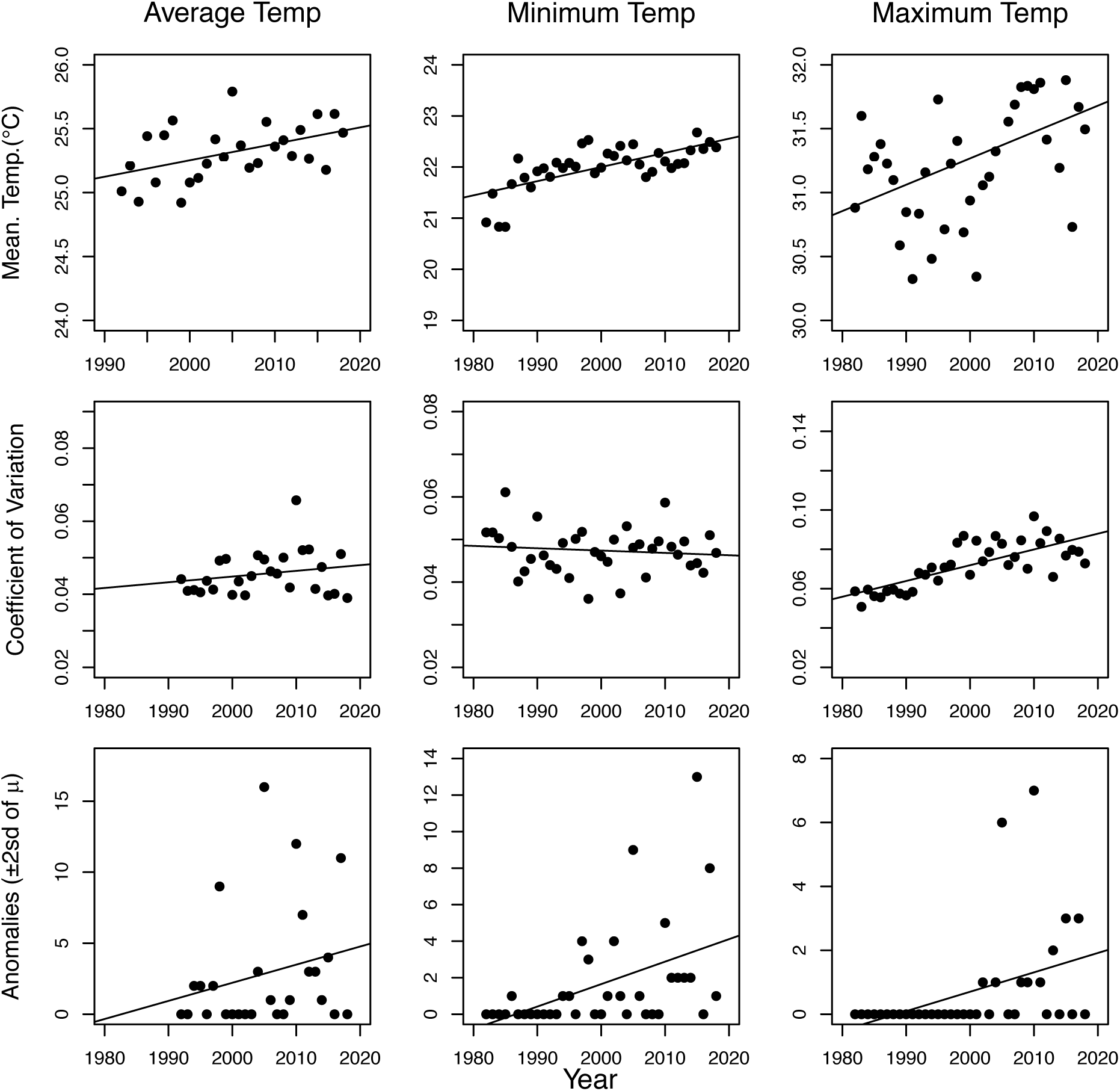
Patterns in temperature variables (average, minimum and maximum daily temperature) at La Selva from 1982-2018. Each point represents a year of data. Graphs in the first row represent annual means of daily values for each temperature variable. The second row displays the intra-annual Coefficient of Variation. The third row displays temperature anomalies measured as the sum of daily values exceeding 2 SD of the annual mean.

**Fig. S8.**
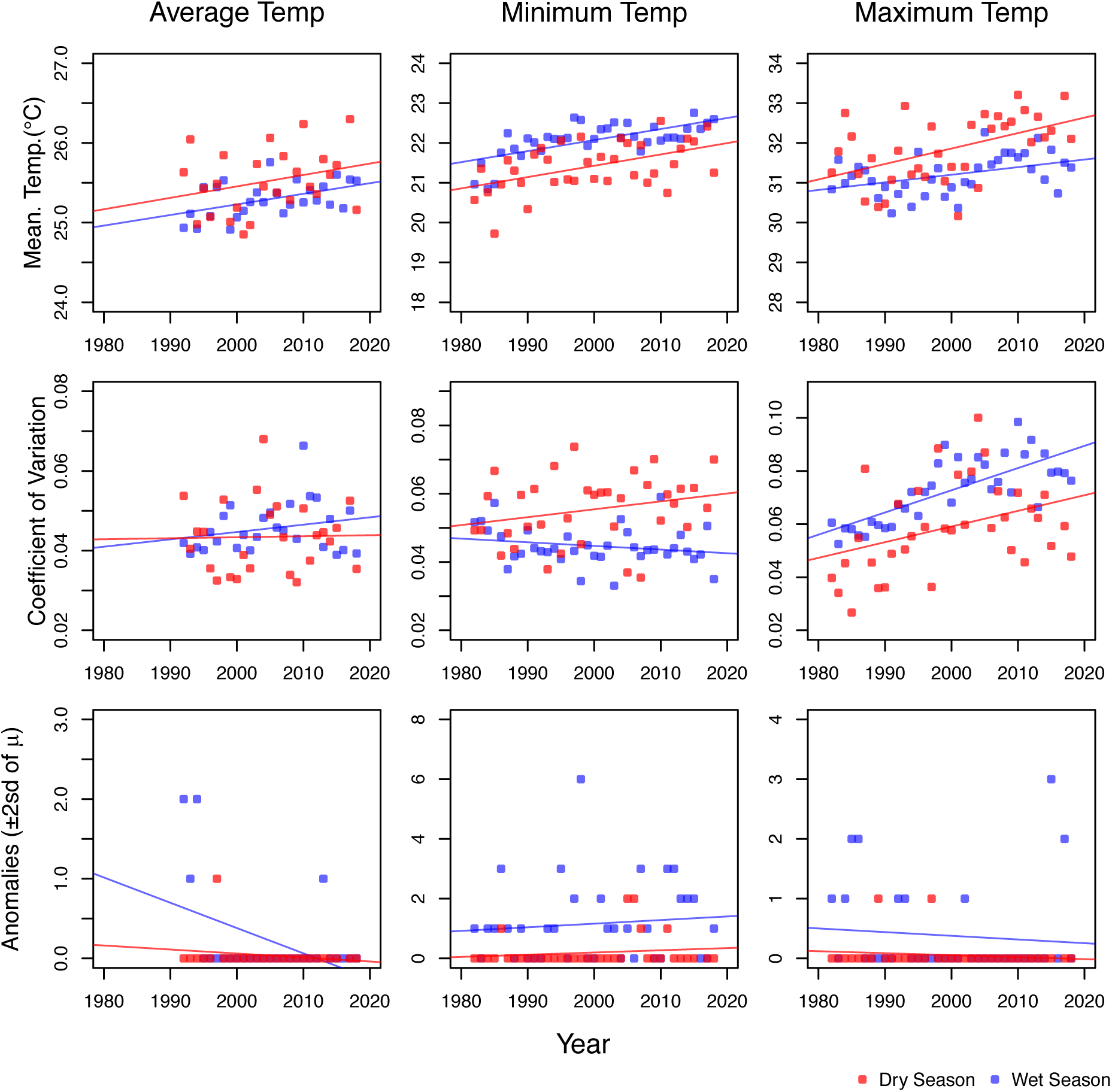
Patterns in temperature variables at La Selva from 1982-2018 for the wet (blue) and dry (red) season. The wet season includes data from May-December and results for the dry season include January-April. Each point represents a year of data. Graphs in the first row represent annual means of daily values for each temperature variable. The second row displays the intra-annual Coefficient of Variation. The third row displays temperature anomalies measured as the sum of daily values exceeding 2 SD of the annual mean.

**Fig. S9.**
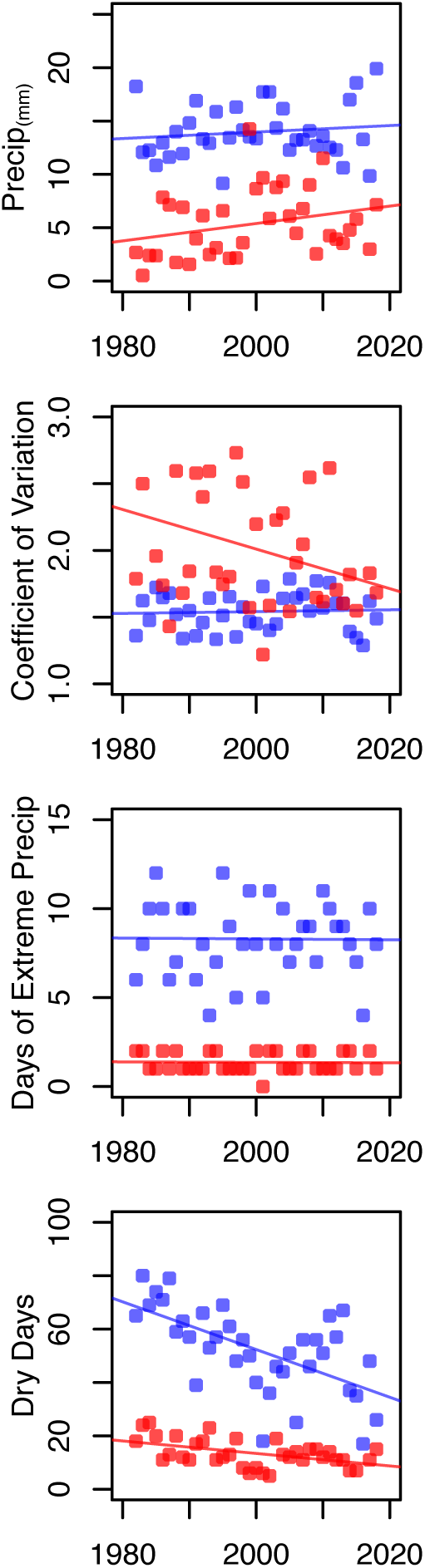
Patterns in precipitation variables at La Selva from 1982-2018 for the wet (blue) and dry (red) season. The wet season includes data from May-December and results for the dry season include January-April. Precip (mm) is calculated as the annual mean of daily precipitation, the Coefficient of Variation (CV) in precipitation is calculated as intra-annual CV, days of extreme precipitation are counts of daily precipitation exceeding 2.5 SD of annual mean precipitation, and dry days are calculated as total days within a year with zero rainfall.

**Fig. S10.**
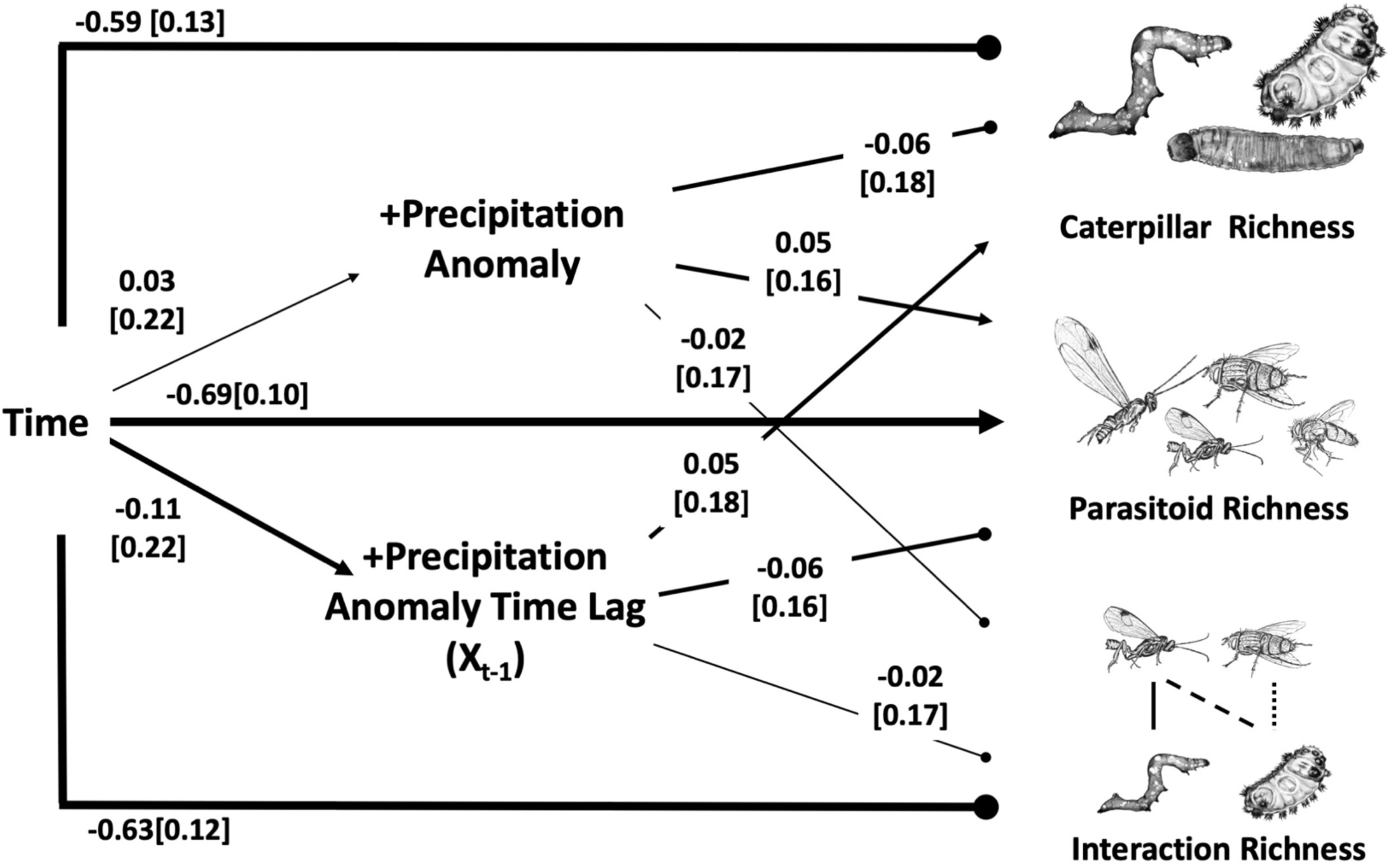
Structural equation models (SEM) testing the effects of positive temperature anomalies and their one year time lag (x_t-1_) on caterpillar, parasitoid and interaction richness. Time is an exogenous variable representing year and the endogenous variables include richness, precipitation anomalies, and time lags; model fit: χ^2^ =0.682, p=0.409, df=1. Path coefficients are standardized and width of arrows are scaled based on magnitude of path coefficients. Arrows represent positive associations and lines with circle represent negative associations. Parasitoid illustrations by M.L.F. Caterpillar images by B.L.

**Fig. S11.**
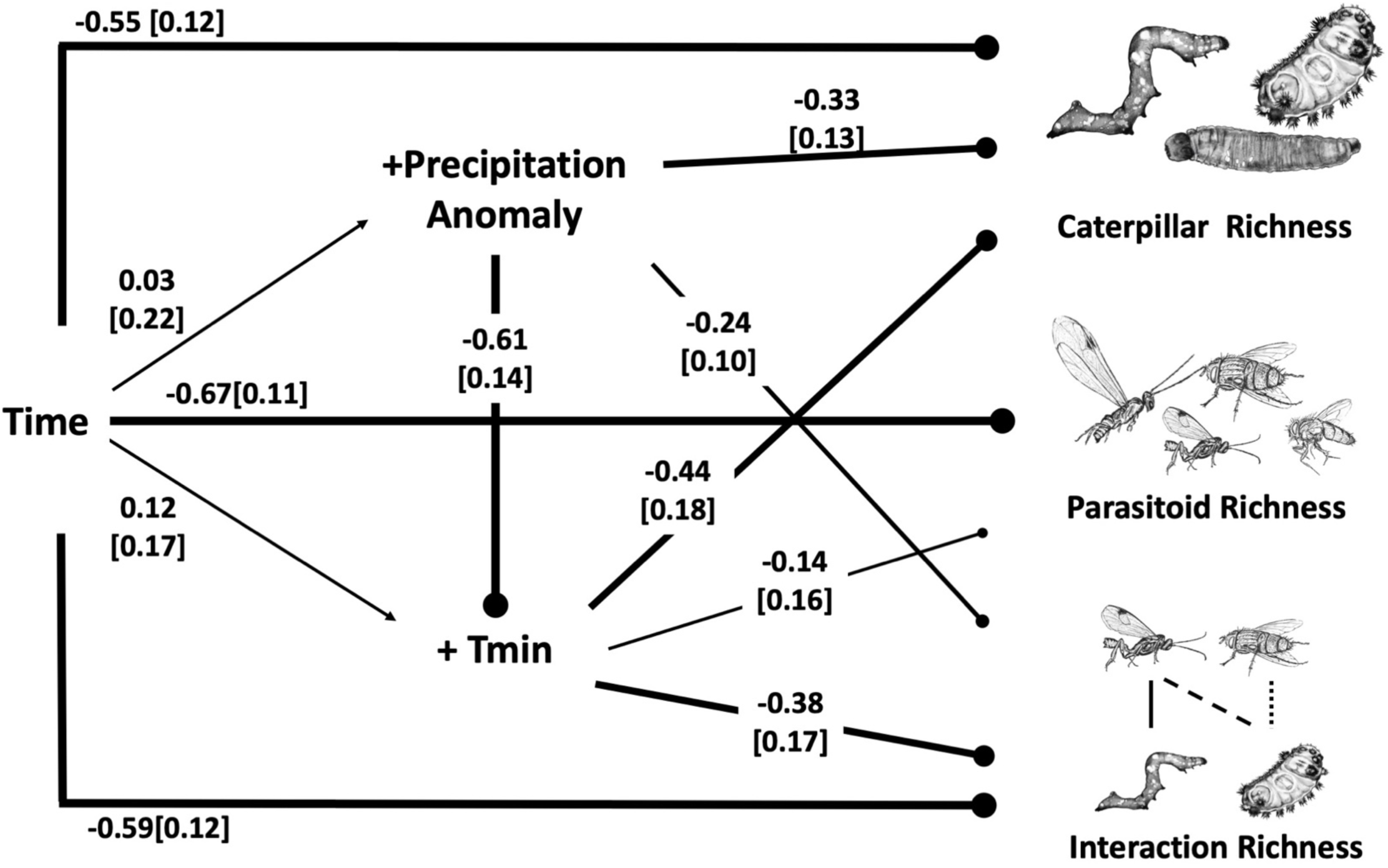
Structural equation models (SEM) testing the effects of minimum temperature (Tmin), and precipitation anomalies on caterpillar, parasitoid and interaction richness. Time is an exogenous variable representing year, and the endogenous variables include richness, Tmin, and precipitation anomalies; <u>m</u>odel fit: *χ*^2^ =0.02, p=0.89, df=1. Path coefficients are standardized and width of arrows are scaled based on magnitude of path coefficients. Arrows represent positive associations and lines with circle represent negative associations. Parasitoid illustrations by M.L.F. Caterpillar images by B.L.

**Table S1.**
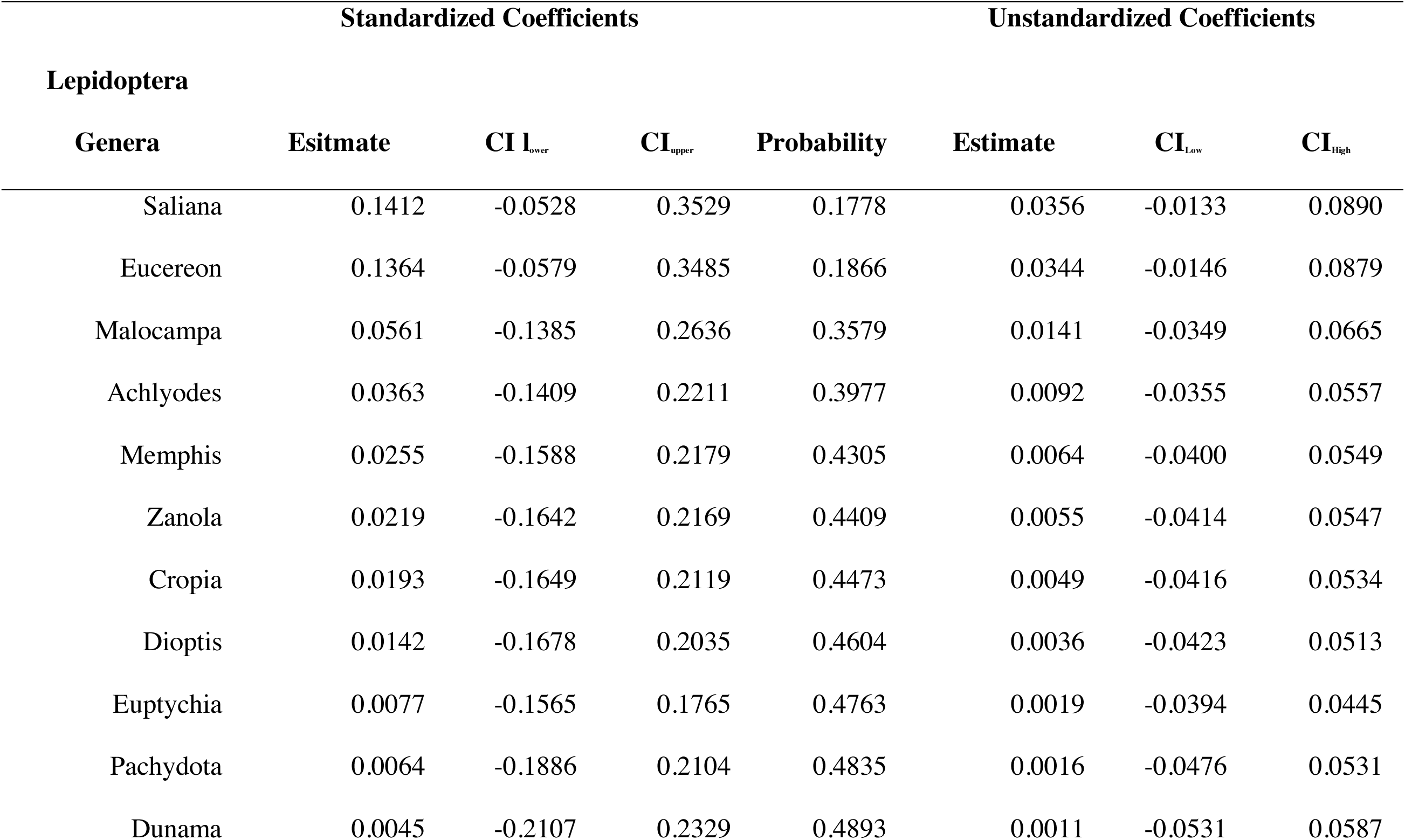
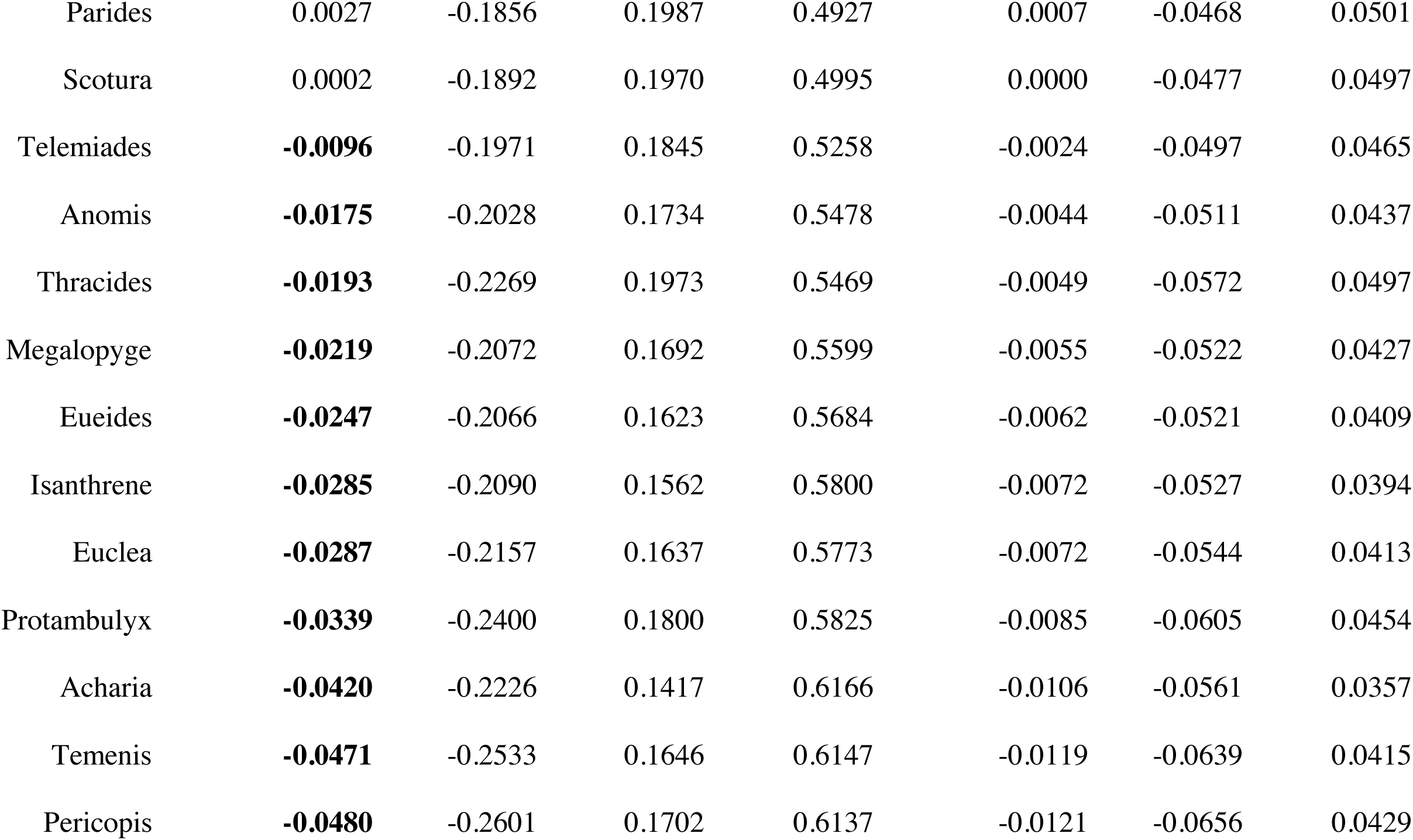
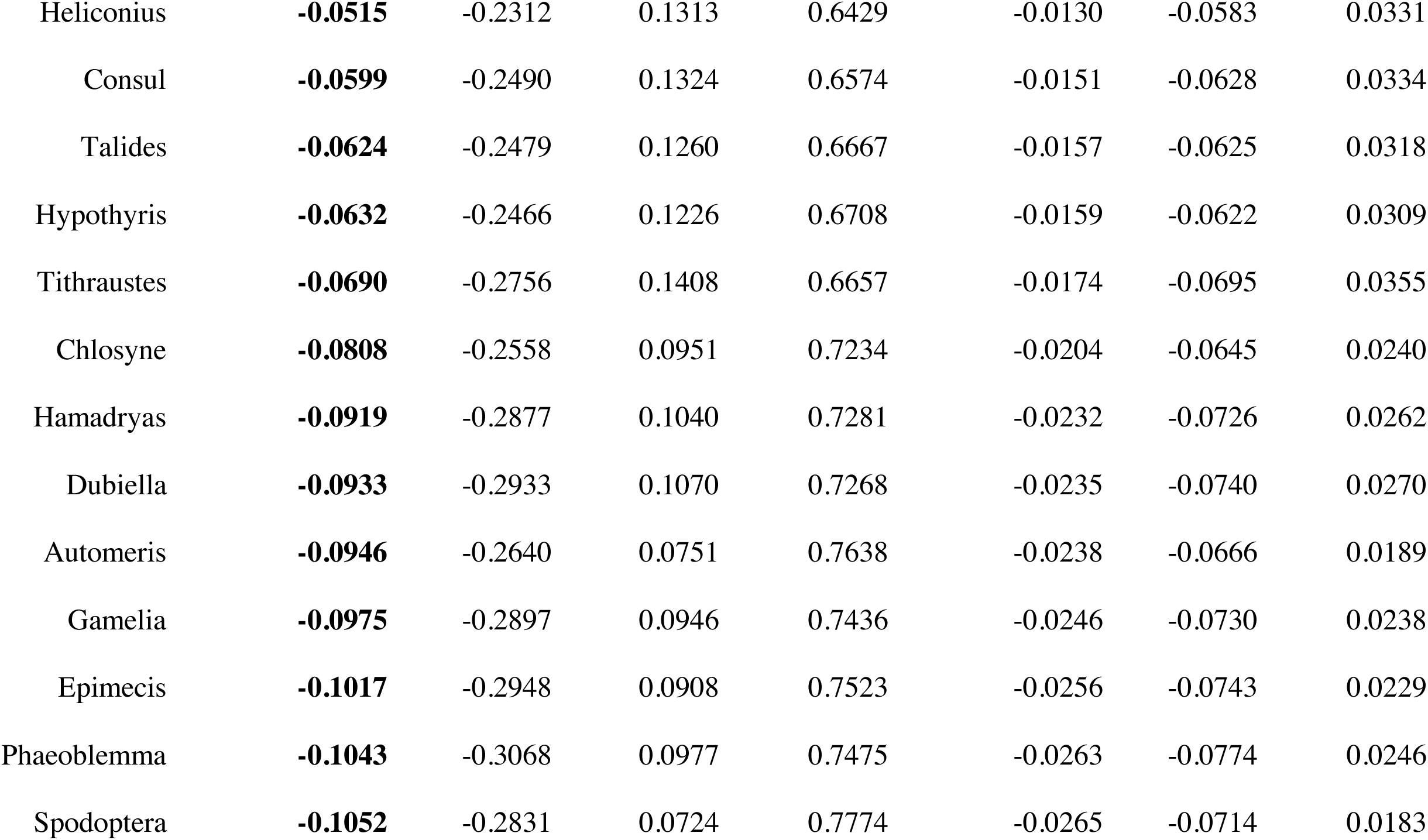
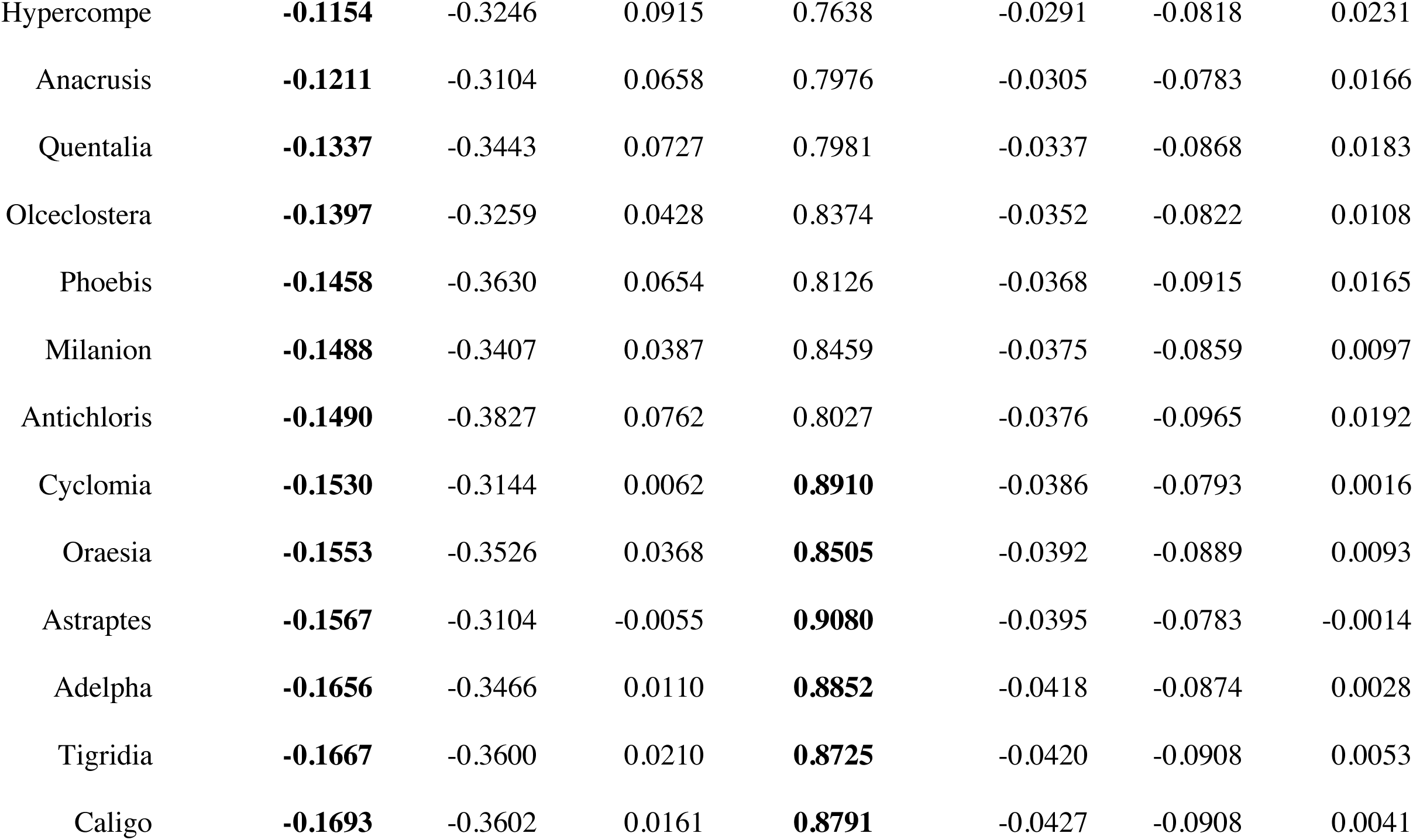
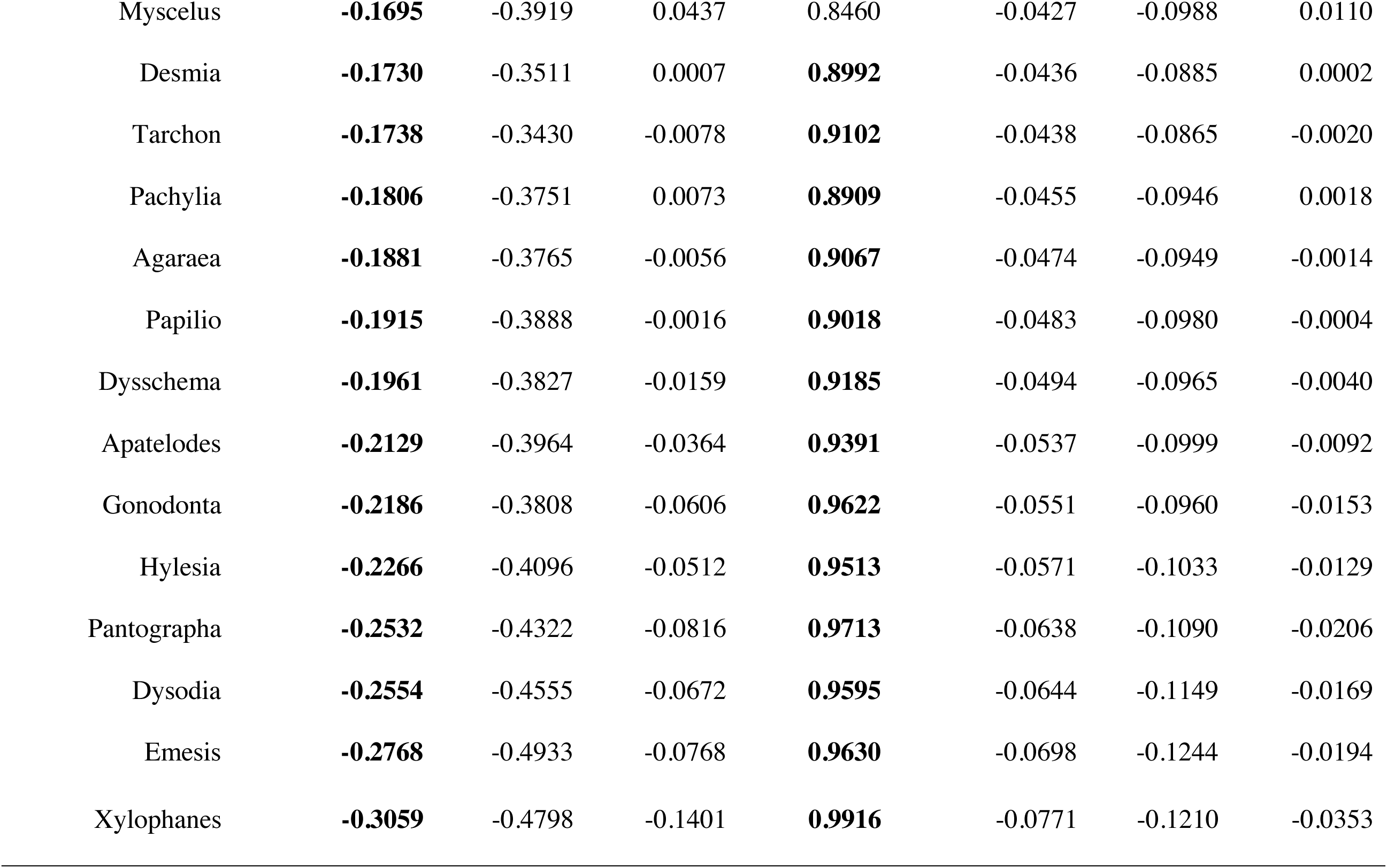
Estimates of standardized and unstandardized beta coefficients and associated 80% credible intervals for each Lepidoptera genera nested in a hierarchical Bayesian model that modeled frequency across years.

**Table S2.**
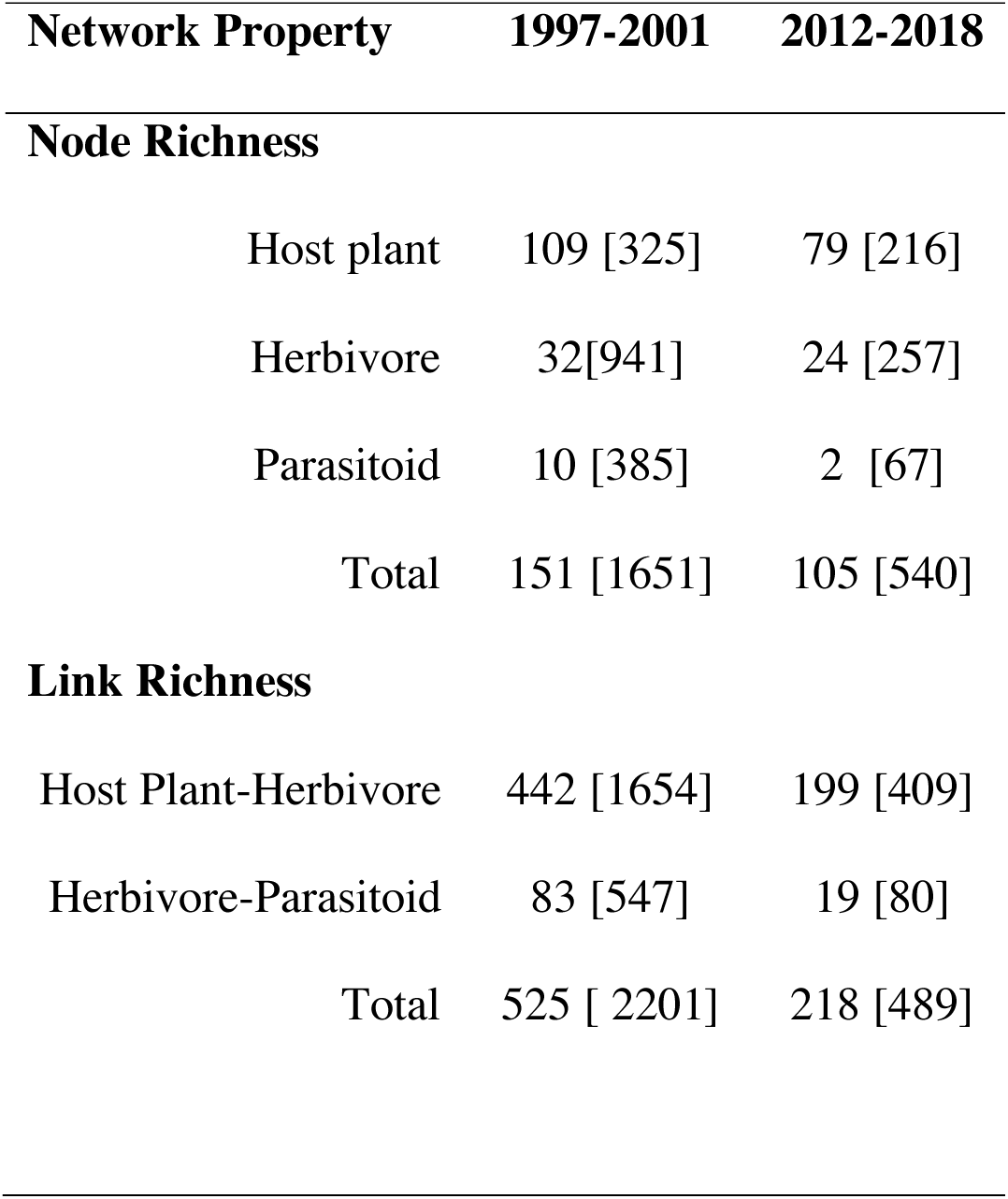
Quantitative comparisons among observed network metrics summed for the first (1997-2001) and last (2012-2018) five years of data. Values represent network properties summarized at the level of taxonomic families, and (in square brackets) at the level of species.

**Table S3.**
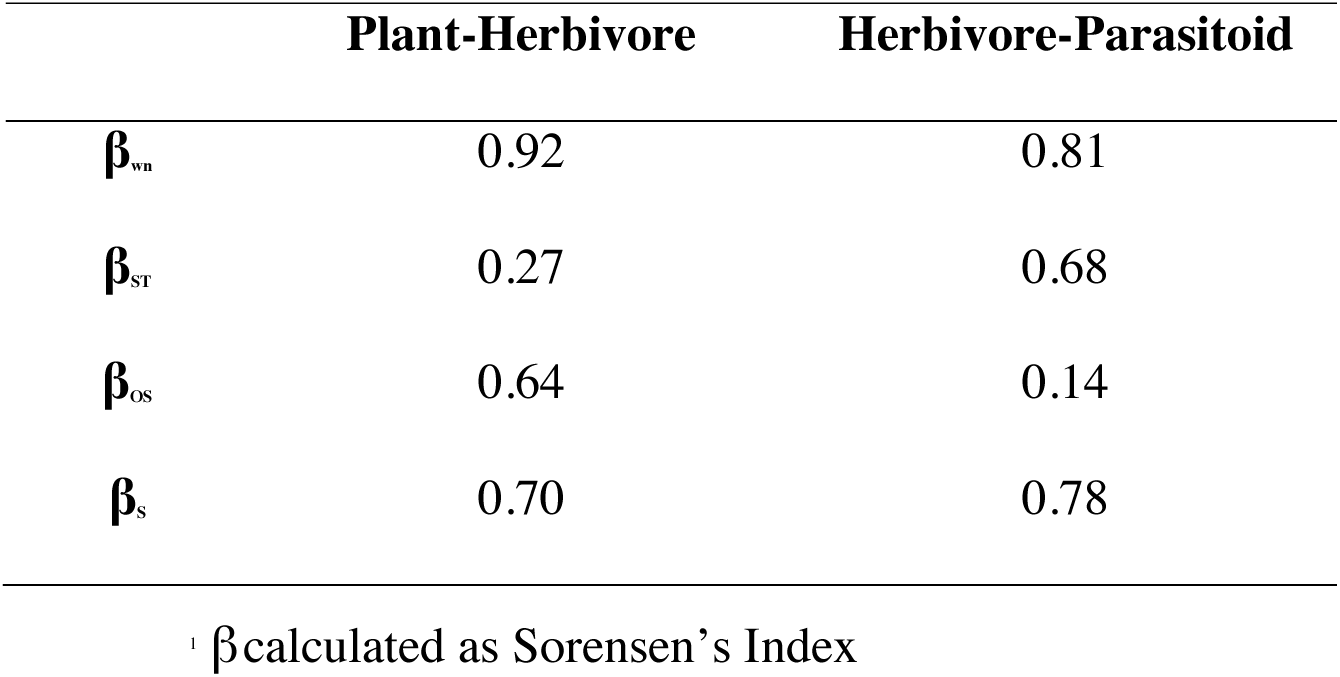
Interaction turnover (β_wn_) and its components among plant-herbivore and herbivore-parasitoid species-level interactions for networks representing the first (1997-2001) and last (2012-2018) five years of data. Interaction turnover among two networks is the sum of turnover owed to differences in species composition (β_ST_) and shared species interacting differently (β_OS_). Species turnover (β_S_) is included for reference.

|  | **Plant-Herbivore** | **Herbivore-Parasitoid** |
| --- | --- | --- |
| $\beta_{wn}$ | 0.92 | 0.81 |
| $\beta_{st}$ | 0.27 | 0.68 |
| $\beta_{os}$ | 0.64 | 0.14 |
| $\beta_s$ | 0.70 | 0.78 |

<sup>1</sup> $\beta$ calculated as Sorensen's Index

**Table S4.**
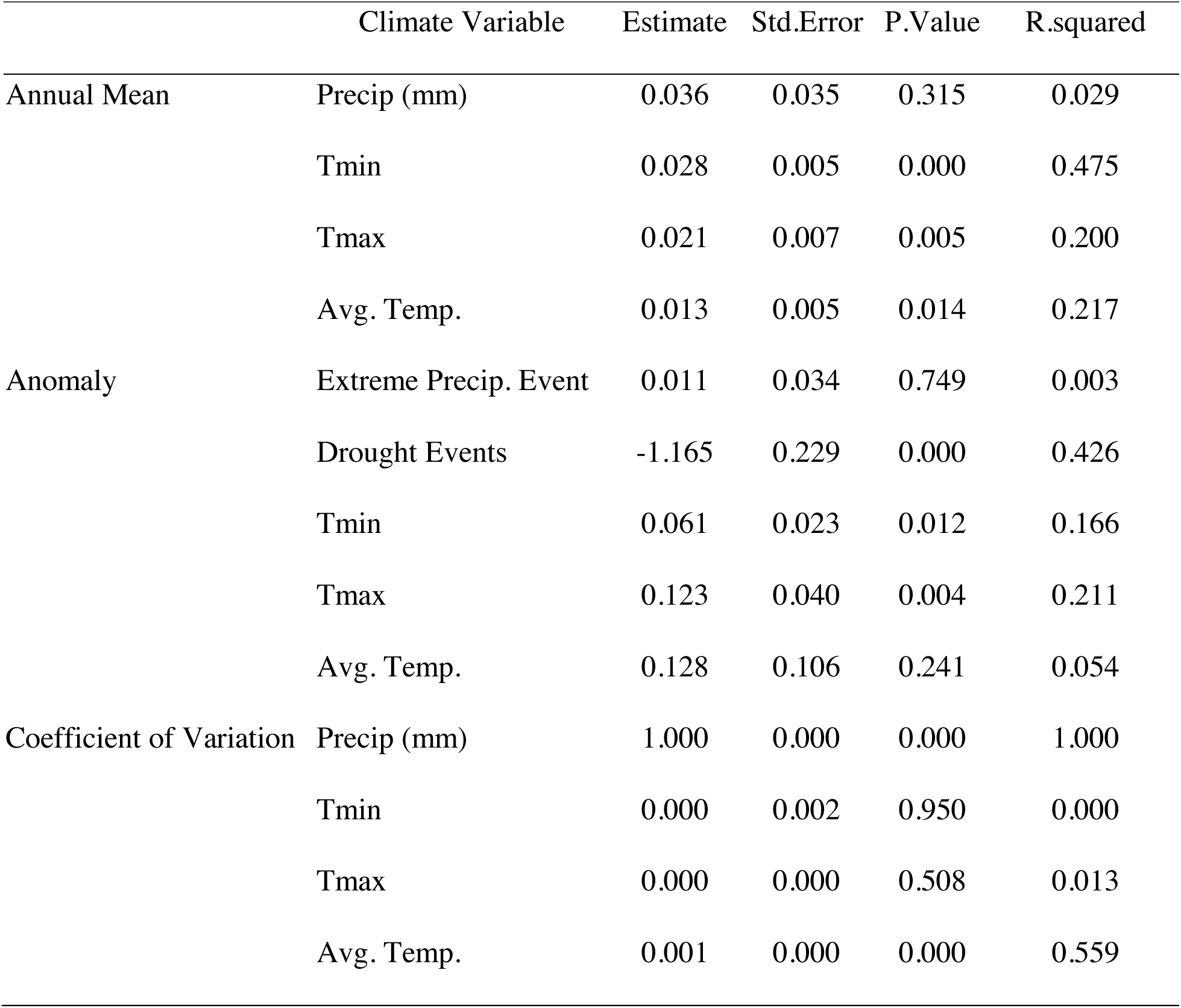
Linear model estimates and fit for various climate variables collected from 1983-2018.

**Table S5.**
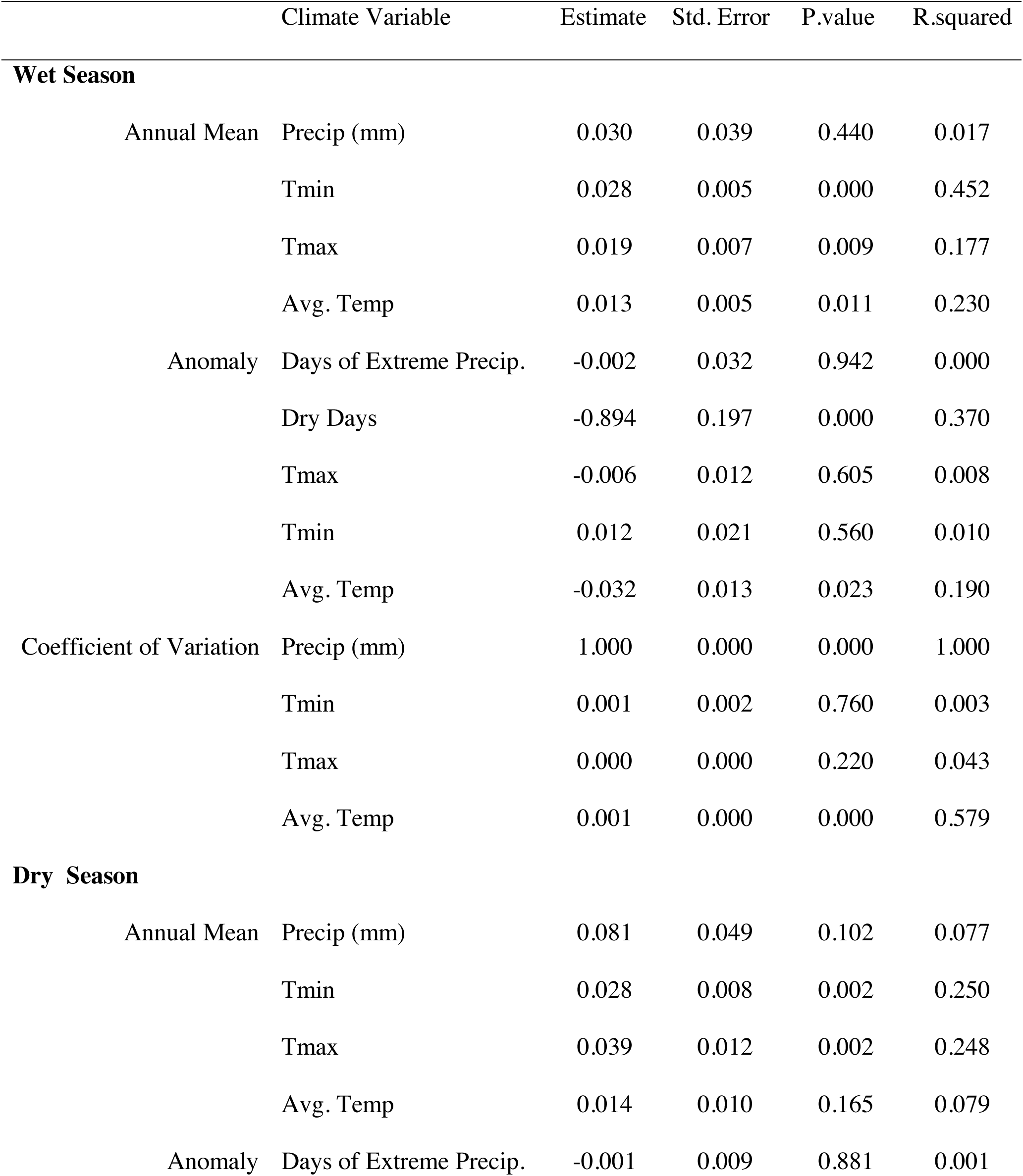
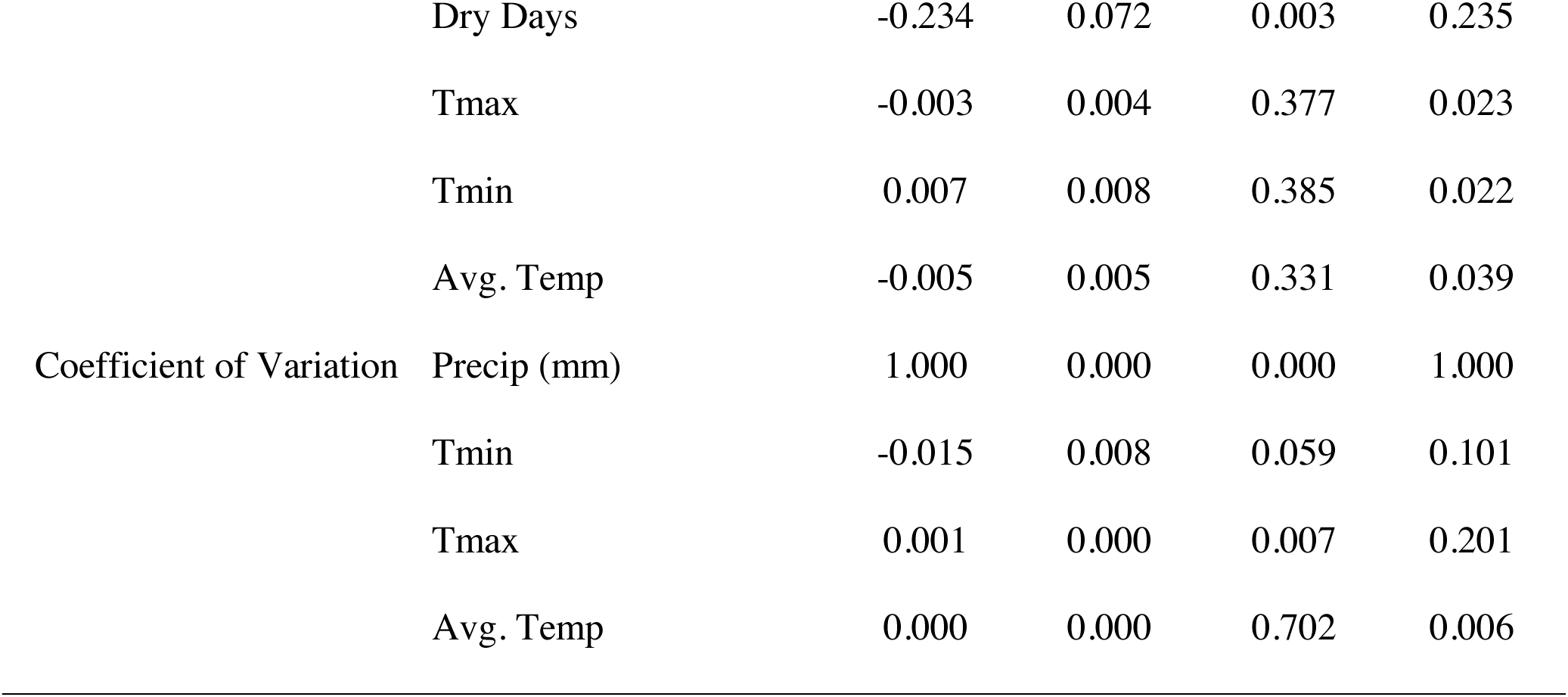
Linear model estimates and fit for various climate variables collected from 1983-2018. Data is displayed for wet (May-December) and dry(January-April) seasons separately.

**Table S6.**
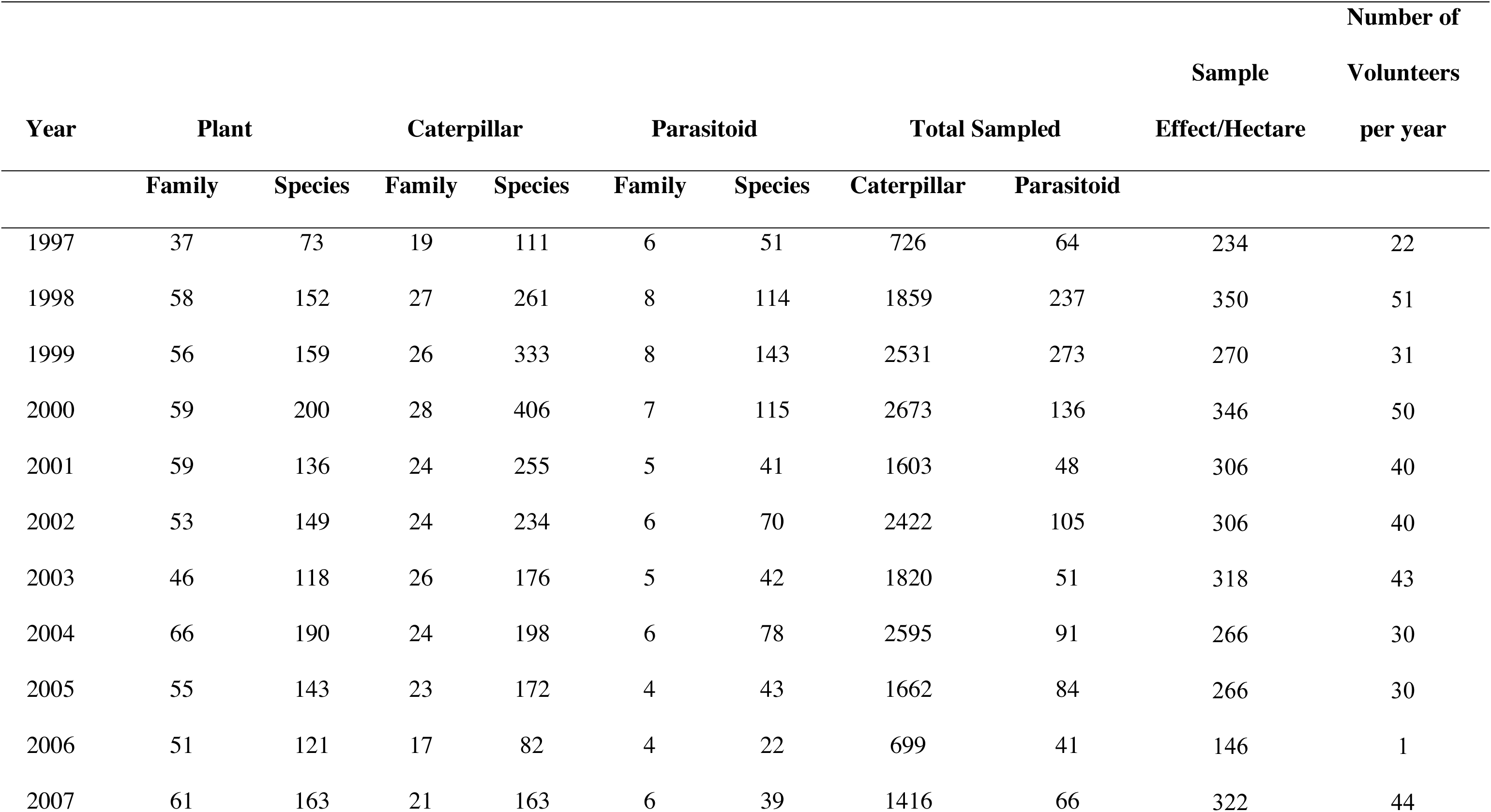
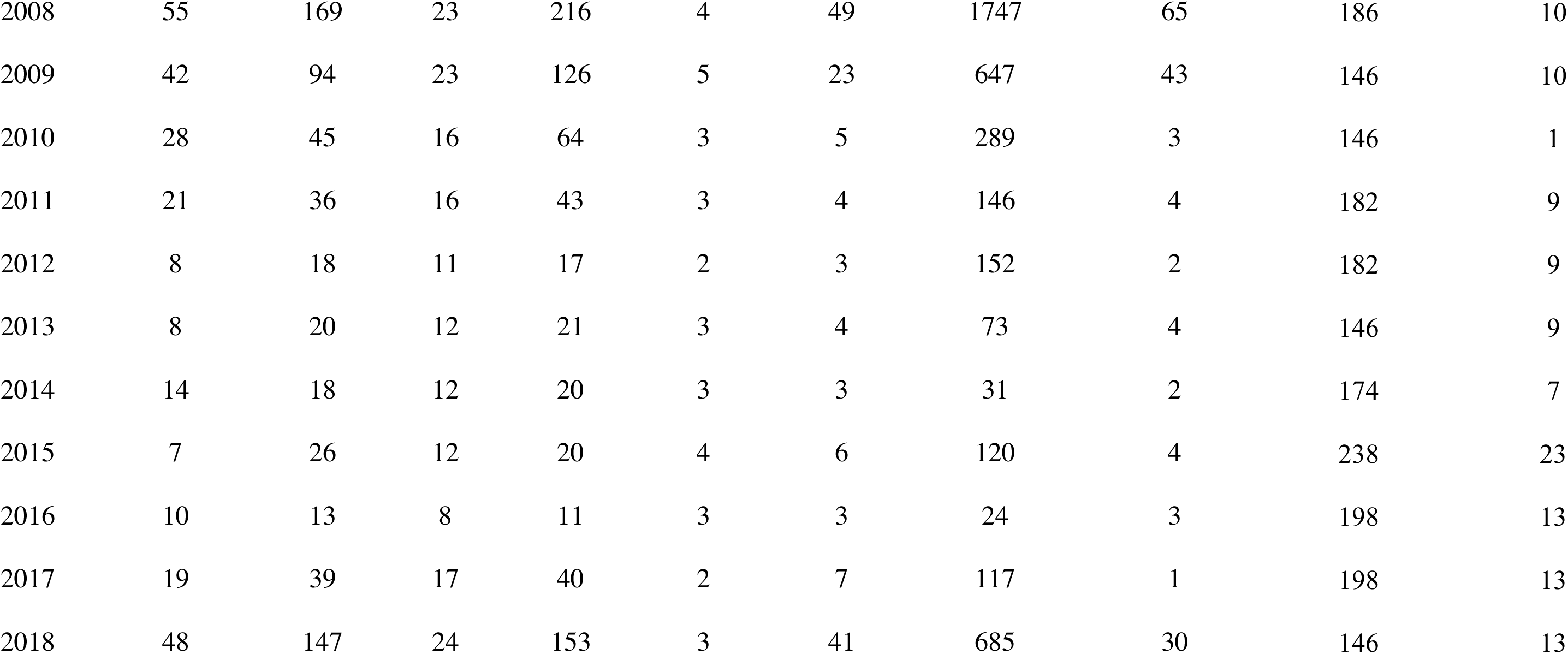
Summary statistics describing annual sampling totals across plant, caterpillar and parasitoid taxonomical groupings.

