## Supplementary Material for "Loss of dominant caterpillar genera in a protected tropical forest"

**Table S3.** Interaction turnover ($\beta_{wn}$) and its components among plant-herbivore and herbivore-parasitoid species-level interactions for networks representing the first (1997-2001) and last (2012-2018) five years of data. Interaction turnover among two networks is the sum of turnover owed to differences in species composition ($\beta_{ST}$) and shared species interacting differently ($\beta_{OS}$). Species turnover ($\beta_{S}$) is included for reference.

**Table S4.** Linear model estimates and fit for various climate variables collected from 1983-2018.

**Table S5**. Linear model estimates and fit for various climate variables collected from 1983-2018. Data is displayed for wet (May-December) and dry(January-April)  seasons separately.

**Table S6**. Summary statistics describing annual sampling totals across plant, caterpillar and parasitoid taxonomical groupings.

**Figure S1**

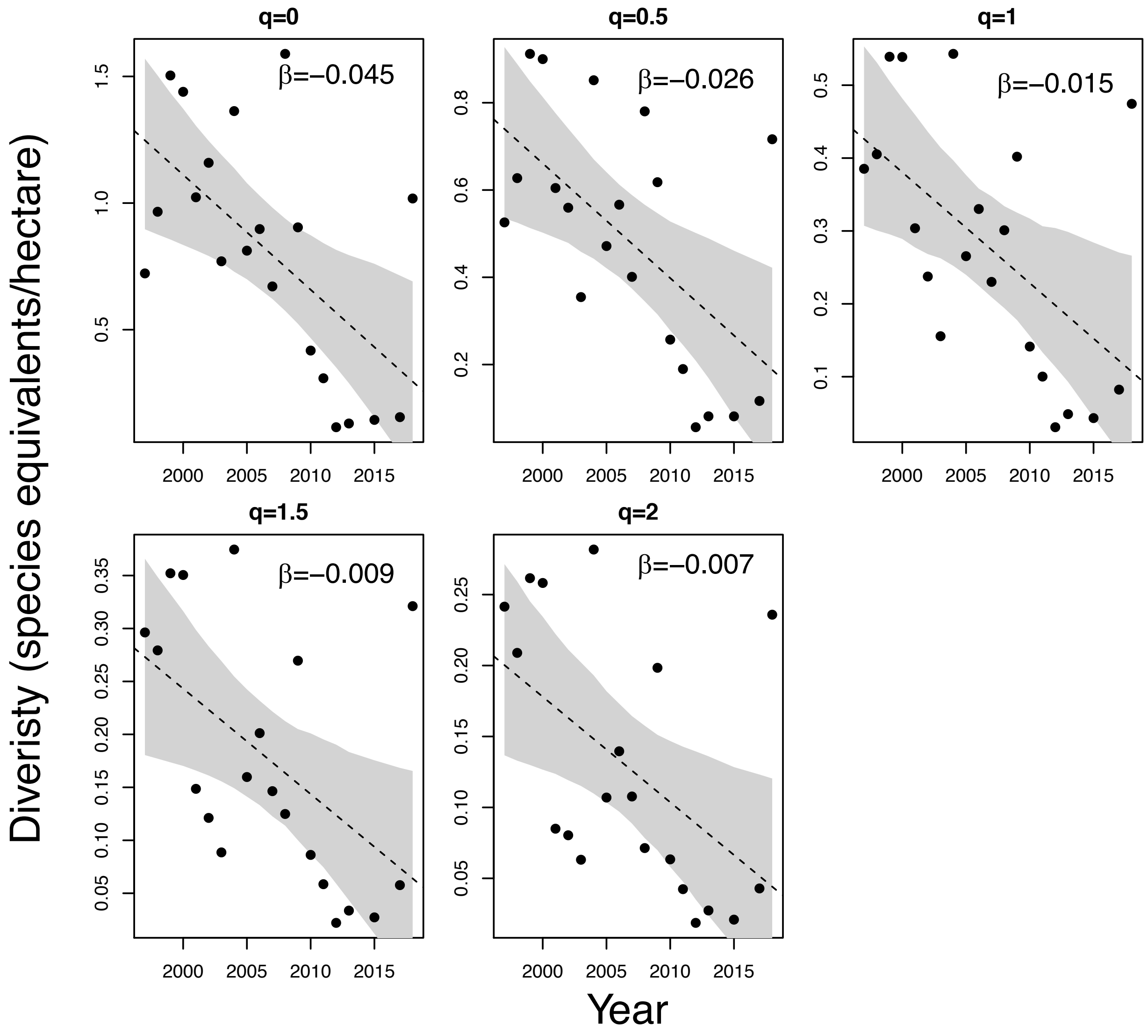
q=0
β=–0.045
q=0.5
β=–0.026
q=1
β=–0.015
q=1.5
β=–0.009
q=2
β=–0.007
Diveristy (species equivalents/hectare)
Year
2000
2005
2010
2015

**Figure S2**

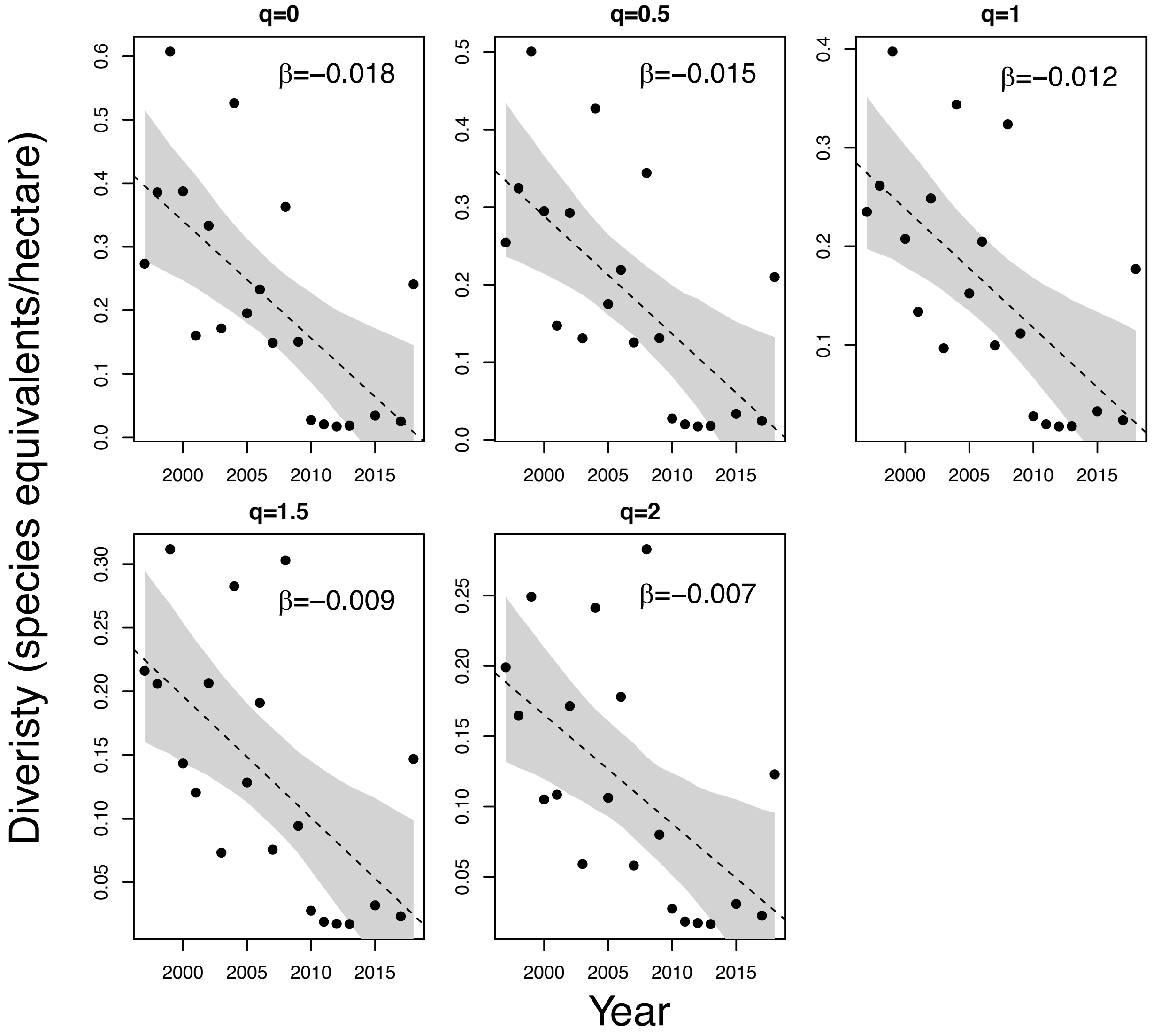
q=0
β=−0.018
q=0.5
β=−0.015
q=1
β=−0.012
q=1.5
β=−0.009
q=2
β=−0.007
Diveristy (species equivalents/hectare)
Year

**Figure S3.**

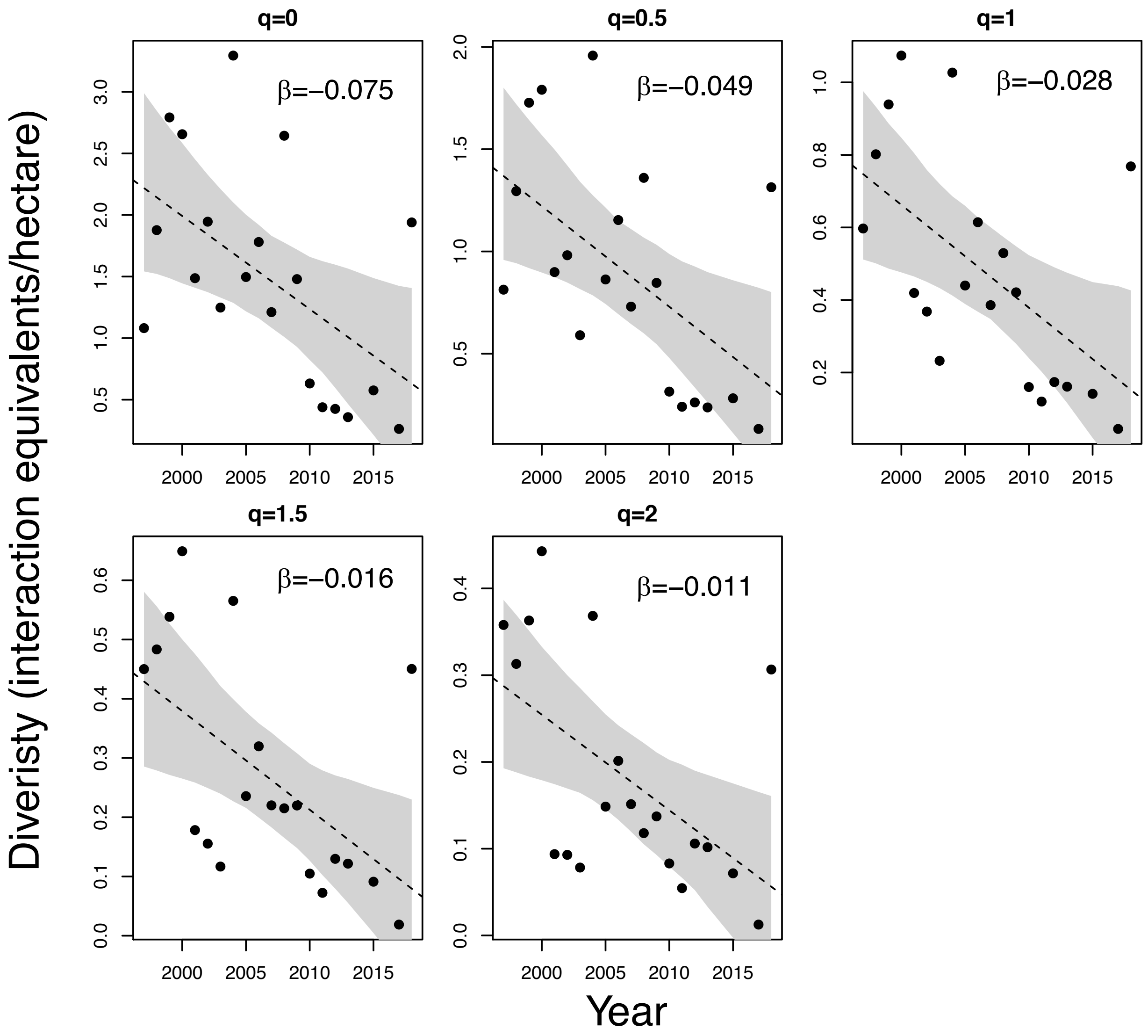

q=0
β=−0.075
q=0.5
β=−0.049
q=1
β=−0.028
q=1.5
β=−0.016
q=2
β=−0.011
Diveristy (interaction equivalents/hectare)
Year

**Figure S4.**

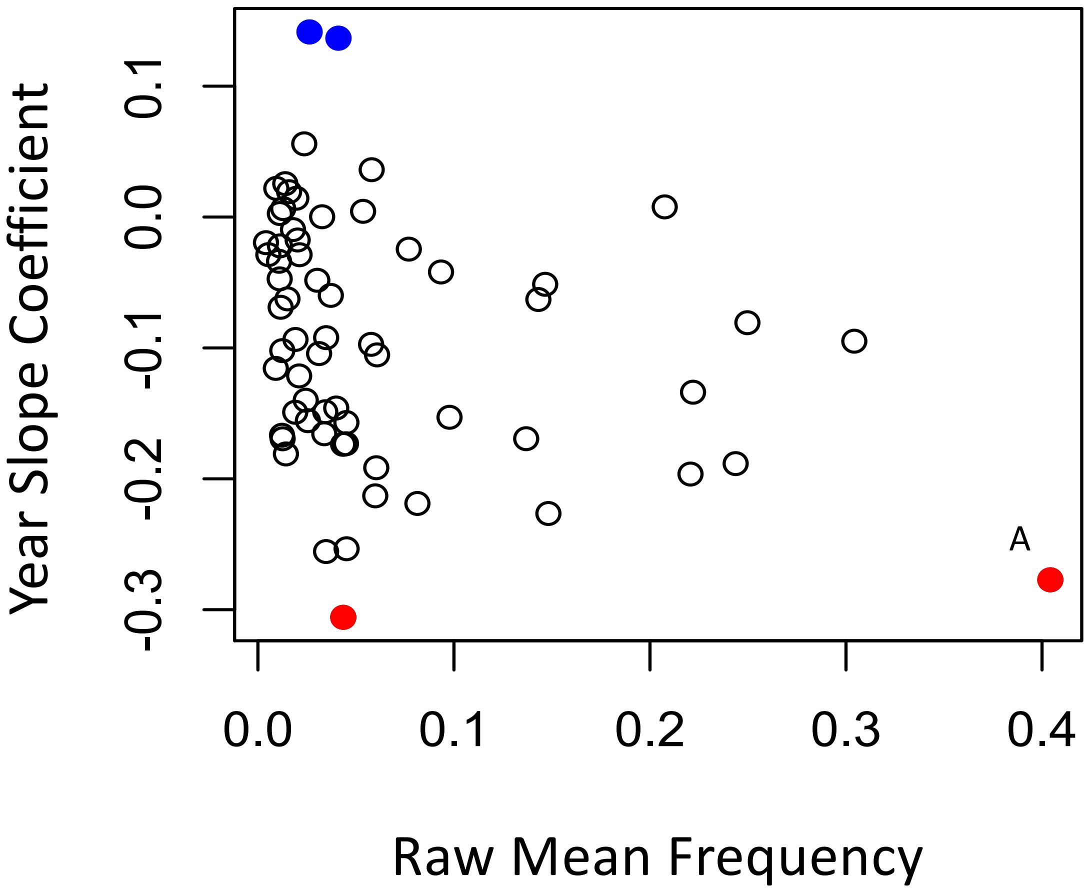
Year Slope Coefficient
0.1
0.0
-0.1
-0.2
-0.3
A
0.0
0.1
0.2
0.3
0.4
Raw Mean Frequency

**Figure S5.**

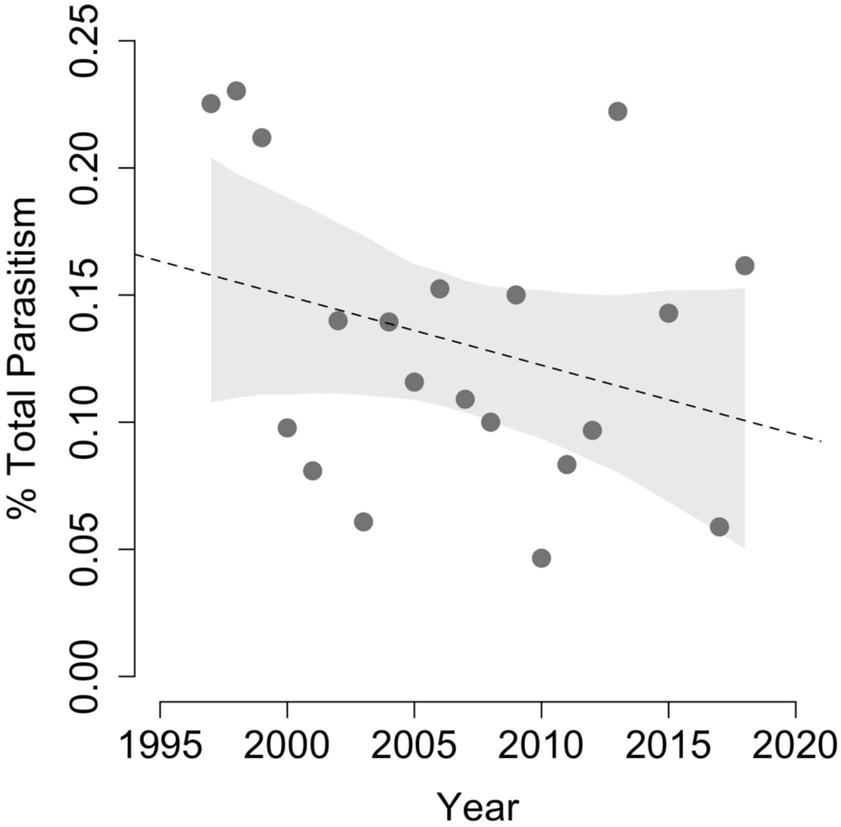

0.25
0.20
0.15
0.10
0.05
0.00
% Total Parasitism
1995
2000
2005
2010
2015
2020
Year

**Figure S6.**

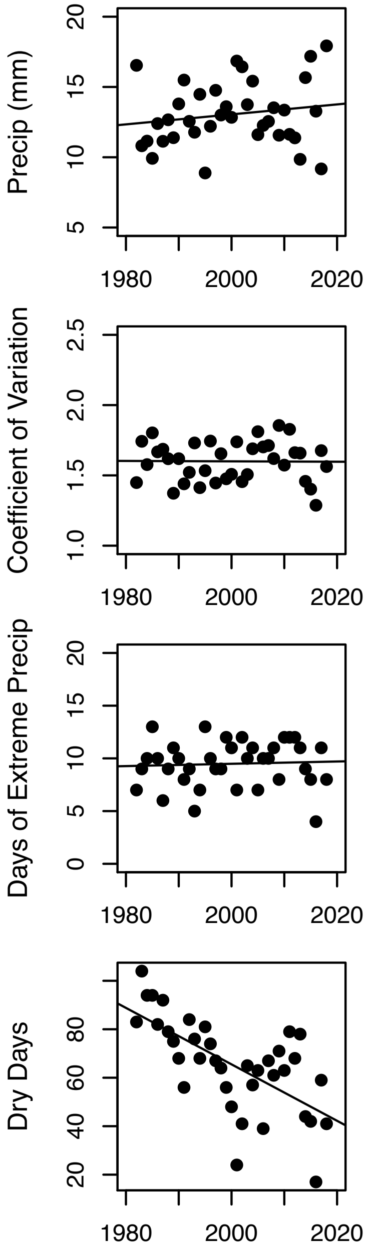

Precip (mm)
5
10
15
20
1980
2000
2020
Coefficient of Variation
1.0
1.5
2.0
2.5
1980
2000
2020
Days of Extreme Precip
0
5
10
15
20
1980
2000
2020
Dry Days
20
40
60
80
1980
2000
2020

**Figure S7.**

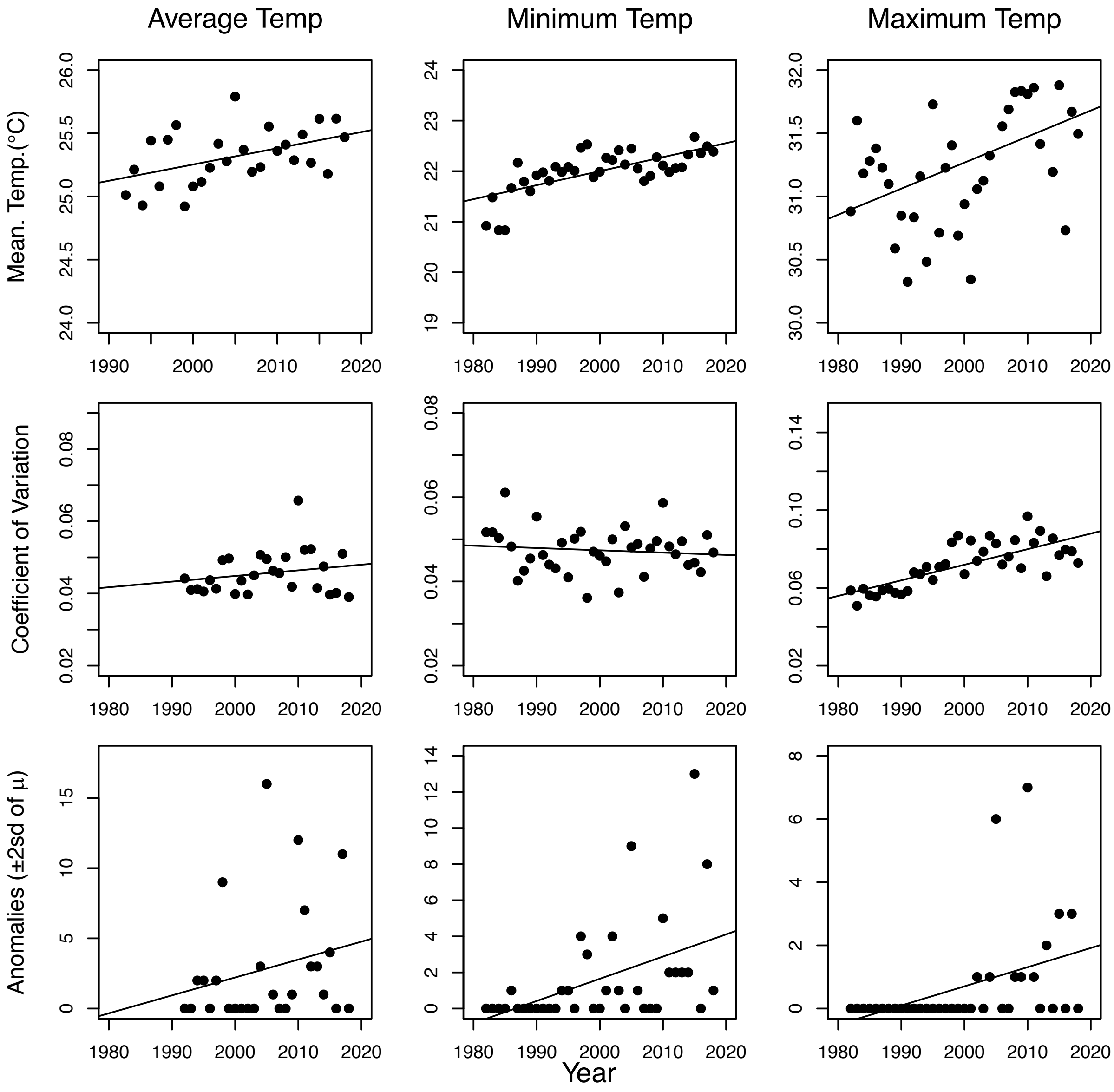

Average Temp
Minimum Temp
Maximum Temp
Mean. Temp.(°C)
Coefficient of Variation
Anomalies (±2sd of μ)
Year

**Figure S8.**

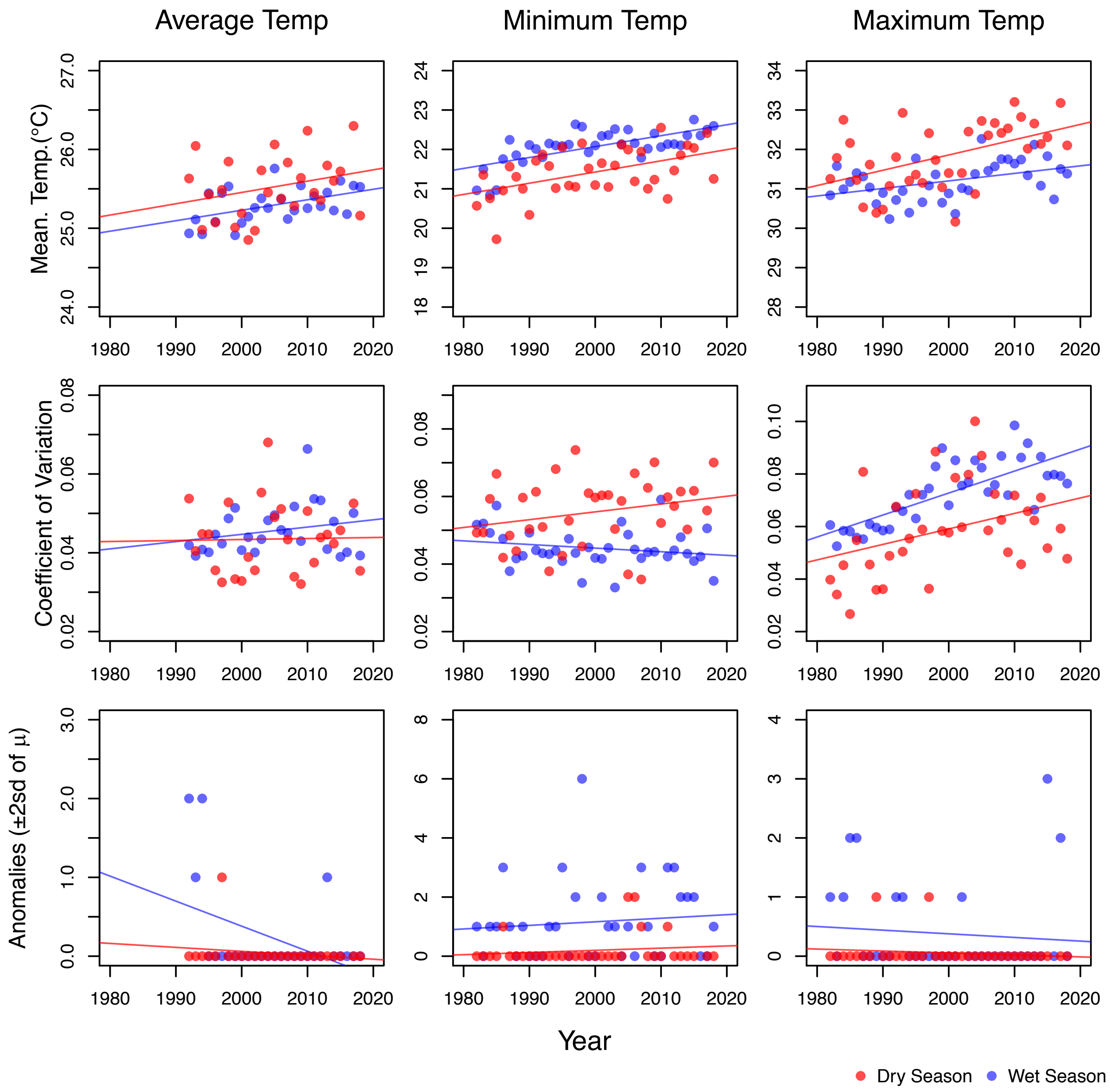

Average Temp
Minimum Temp
Maximum Temp
Mean. Temp.(°C)
Coefficient of Variation
Anomalies (±2sd of μ)
Year
Dry Season
Wet Season

**Figure S9.**

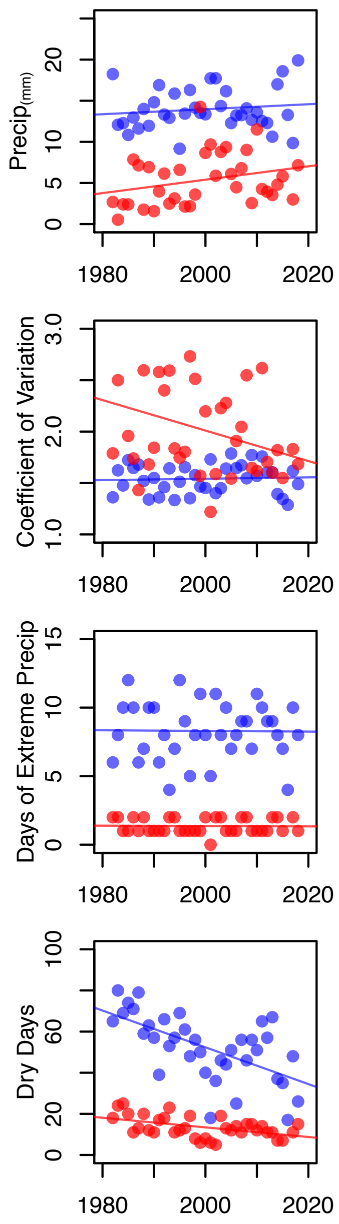

Precip(mm)
0
5
10
20
1980
2000
2020
Coefficient of Variation
1.0
2.0
3.0
1980
2000
2020
Days of Extreme Precip
0
5
10
15
1980
2000
2020
Dry Days
0
20
60
100
1980
2000
2020

**Figure S10.**

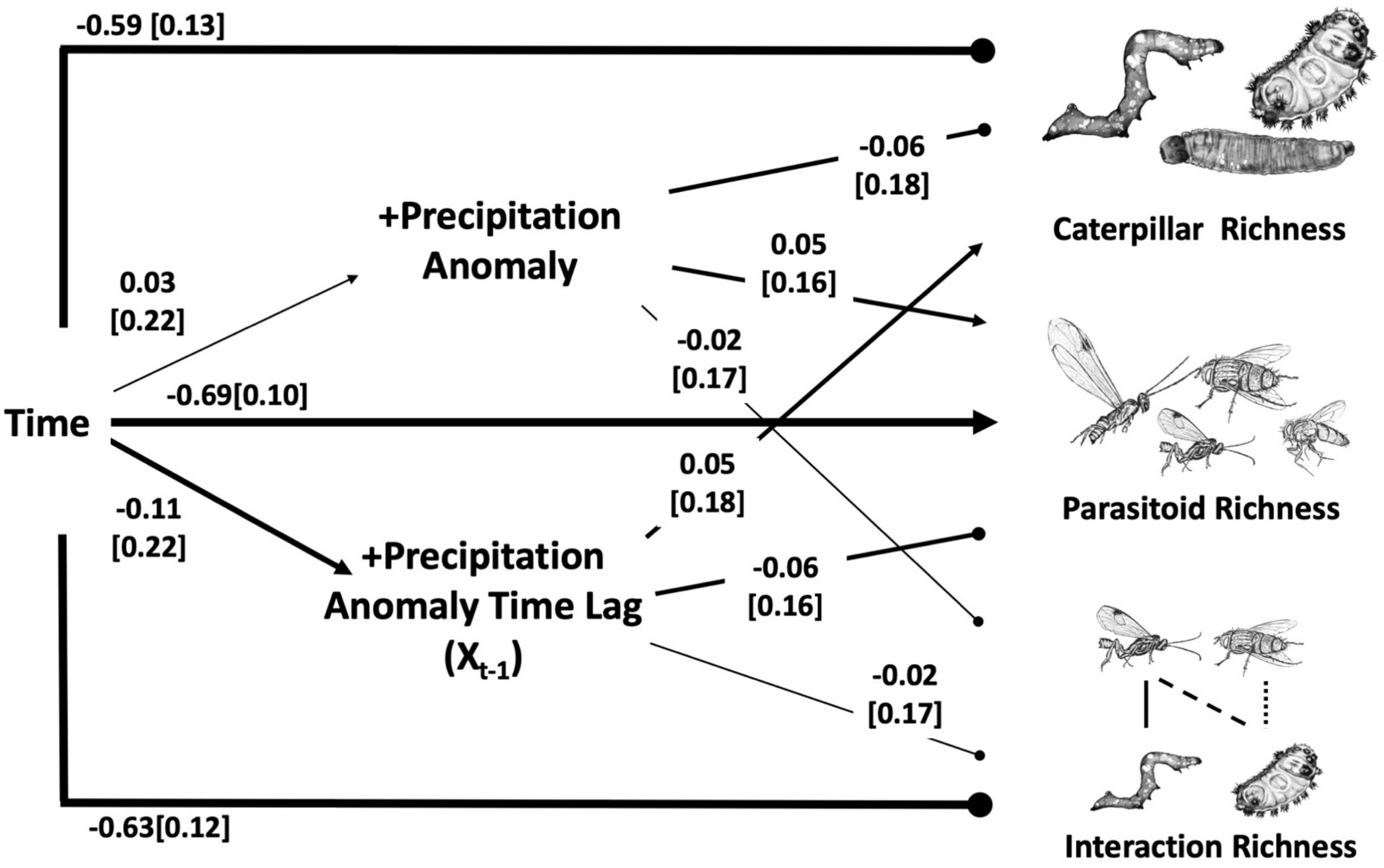

-0.59 [0.13]
-0.06
[0.18]
+Precipitation
Anomaly
0.03
[0.22]
0.05
[0.16]
-0.02
[0.17]
-0.69[0.10]
Time
0.05
[0.18]
-0.11
[0.22]
+Precipitation
Anomaly Time Lag
($X_{t-1}$)
-0.06
[0.16]
-0.02
[0.17]
-0.63[0.12]
Caterpillar Richness
Parasitoid Richness
Interaction Richness

**FigS11**

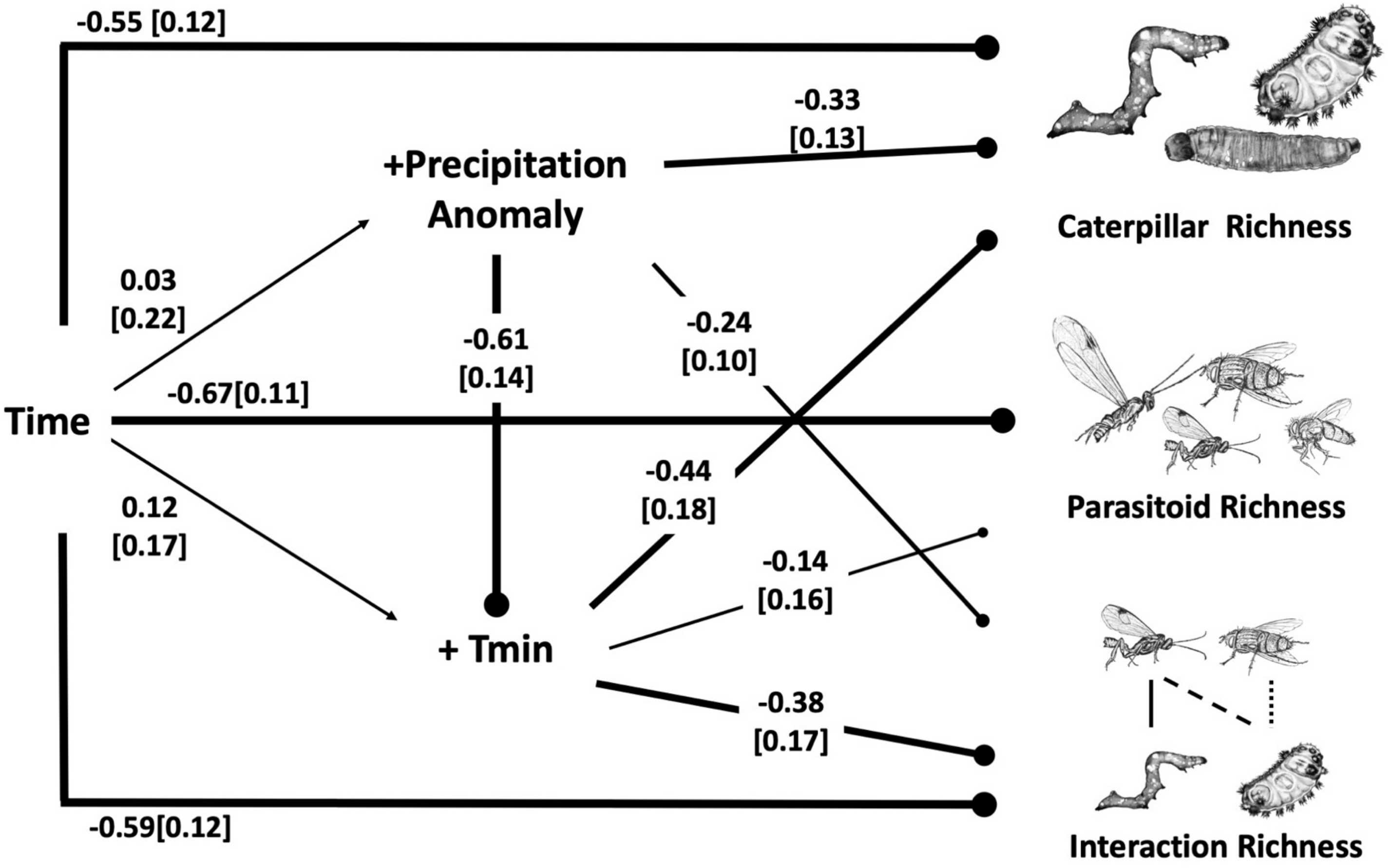
-0.55 [0.12]
-0.33
[0.13]
+Precipitation
Anomaly
Caterpillar Richness
0.03
[0.22]
-0.61
[0.14]
-0.24
[0.10]
Time
-0.67[0.11]
Parasitoid Richness
-0.44
[0.18]
0.12
[0.17]
-0.14
[0.16]
+ Tmin
-0.38
[0.17]
-0.59[0.12]
Interaction Richness

**Table S1**

| | Standardized Coefficients | | | | Unstandardized Coefficients | | |
|---|---|---|---|---|---|---|---|
| **Lepidoptera** | | | | | | | |
| **Genera** | **Esitmate** | **CI $l_{ower}$** | **$CI_{upper}$** | **Probability** | **Estimate** | **$CI_{Low}$** | **$CI_{High}$** |
| Saliana | 0.1412 | -0.0528 | 0.3529 | 0.1778 | 0.0356 | -0.0133 | 0.0890 |
| Eucereon | 0.1364 | -0.0579 | 0.3485 | 0.1866 | 0.0344 | -0.0146 | 0.0879 |
| Malocampa | 0.0561 | -0.1385 | 0.2636 | 0.3579 | 0.0141 | -0.0349 | 0.0665 |
| Achlyodes | 0.0363 | -0.1409 | 0.2211 | 0.3977 | 0.0092 | -0.0355 | 0.0557 |
| Memphis | 0.0255 | -0.1588 | 0.2179 | 0.4305 | 0.0064 | -0.0400 | 0.0549 |
| Zanola | 0.0219 | -0.1642 | 0.2169 | 0.4409 | 0.0055 | -0.0414 | 0.0547 |
| Cropia | 0.0193 | -0.1649 | 0.2119 | 0.4473 | 0.0049 | -0.0416 | 0.0534 |
| Dioptis | 0.0142 | -0.1678 | 0.2035 | 0.4604 | 0.0036 | -0.0423 | 0.0513 |
| Euptychia | 0.0077 | -0.1565 | 0.1765 | 0.4763 | 0.0019 | -0.0394 | 0.0445 |
| Pachydota | 0.0064 | -0.1886 | 0.2104 | 0.4835 | 0.0016 | -0.0476 | 0.0531 |
| Dunama | 0.0045 | -0.2107 | 0.2329 | 0.4893 | 0.0011 | -0.0531 | 0.0587 |

| | Standardized Coefficients | | | | Unstandardized Coefficients | | |
|---|---|---|---|---|---|---|---|
| **Lepidoptera** | | | | | | | |
| **Genera** | **Esitmate** | **CI l**$_{ower}$ | **CI**$_{upper}$ | **Probability** | **Estimate** | **CI**$_{Low}$ | **CI**$_{High}$ |
| Parides | 0.0027 | -0.1856 | 0.1987 | 0.4927 | 0.0007 | -0.0468 | 0.0501 |
| Scotura | 0.0002 | -0.1892 | 0.1970 | 0.4995 | 0.0000 | -0.0477 | 0.0497 |
| Telemiades | **-0.0096** | -0.1971 | 0.1845 | 0.5258 | -0.0024 | -0.0497 | 0.0465 |
| Anomis | **-0.0175** | -0.2028 | 0.1734 | 0.5478 | -0.0044 | -0.0511 | 0.0437 |
| Thracides | **-0.0193** | -0.2269 | 0.1973 | 0.5469 | -0.0049 | -0.0572 | 0.0497 |
| Megalopyge | **-0.0219** | -0.2072 | 0.1692 | 0.5599 | -0.0055 | -0.0522 | 0.0427 |
| Eueides | **-0.0247** | -0.2066 | 0.1623 | 0.5684 | -0.0062 | -0.0521 | 0.0409 |
| Isanthrene | **-0.0285** | -0.2090 | 0.1562 | 0.5800 | -0.0072 | -0.0527 | 0.0394 |
| Euclea | **-0.0287** | -0.2157 | 0.1637 | 0.5773 | -0.0072 | -0.0544 | 0.0413 |
| Protambulyx | **-0.0339** | -0.2400 | 0.1800 | 0.5825 | -0.0085 | -0.0605 | 0.0454 |
| Acharia | **-0.0420** | -0.2226 | 0.1417 | 0.6166 | -0.0106 | -0.0561 | 0.0357 |
| Temenis | **-0.0471** | -0.2533 | 0.1646 | 0.6147 | -0.0119 | -0.0639 | 0.0415 |
| Pericopis | **-0.0480** | -0.2601 | 0.1702 | 0.6137 | -0.0121 | -0.0656 | 0.0429 |

| | Standardized Coefficients | | | | Unstandardized Coefficients | | |
|---|---|---|---|---|---|---|---|
| **Lepidoptera** | | | | | | | |
| **Genera** | **Esitmate** | **CI $l_{ower}$** | **$CI_{upper}$** | **Probability** | **Estimate** | **$CI_{Low}$** | **$CI_{High}$** |
| Heliconius | **-0.0515** | -0.2312 | 0.1313 | 0.6429 | -0.0130 | -0.0583 | 0.0331 |
| Consul | **-0.0599** | -0.2490 | 0.1324 | 0.6574 | -0.0151 | -0.0628 | 0.0334 |
| Talides | **-0.0624** | -0.2479 | 0.1260 | 0.6667 | -0.0157 | -0.0625 | 0.0318 |
| Hypothyris | **-0.0632** | -0.2466 | 0.1226 | 0.6708 | -0.0159 | -0.0622 | 0.0309 |
| Tithraustes | **-0.0690** | -0.2756 | 0.1408 | 0.6657 | -0.0174 | -0.0695 | 0.0355 |
| Chlosyne | **-0.0808** | -0.2558 | 0.0951 | 0.7234 | -0.0204 | -0.0645 | 0.0240 |
| Hamadryas | **-0.0919** | -0.2877 | 0.1040 | 0.7281 | -0.0232 | -0.0726 | 0.0262 |
| Dubiella | **-0.0933** | -0.2933 | 0.1070 | 0.7268 | -0.0235 | -0.0740 | 0.0270 |
| Automeris | **-0.0946** | -0.2640 | 0.0751 | 0.7638 | -0.0238 | -0.0666 | 0.0189 |
| Gamelia | **-0.0975** | -0.2897 | 0.0946 | 0.7436 | -0.0246 | -0.0730 | 0.0238 |
| Epimecis | **-0.1017** | -0.2948 | 0.0908 | 0.7523 | -0.0256 | -0.0743 | 0.0229 |
| Phaeoblemma | **-0.1043** | -0.3068 | 0.0977 | 0.7475 | -0.0263 | -0.0774 | 0.0246 |
| Spodoptera | **-0.1052** | -0.2831 | 0.0724 | 0.7774 | -0.0265 | -0.0714 | 0.0183 |

| | Standardized Coefficients | | | | Unstandardized Coefficients | | |
|---|---|---|---|---|---|---|---|
| **Lepidoptera** | | | | | | | |
| **Genera** | **Esitmate** | **CI l$_{ower}$** | **CI$_{upper}$** | **Probability** | **Estimate** | **CI$_{Low}$** | **CI$_{High}$** |
| Hypercompe | **-0.1154** | -0.3246 | 0.0915 | 0.7638 | -0.0291 | -0.0818 | 0.0231 |
| Anacrusis | **-0.1211** | -0.3104 | 0.0658 | 0.7976 | -0.0305 | -0.0783 | 0.0166 |
| Quentalia | **-0.1337** | -0.3443 | 0.0727 | 0.7981 | -0.0337 | -0.0868 | 0.0183 |
| Olceclostera | **-0.1397** | -0.3259 | 0.0428 | 0.8374 | -0.0352 | -0.0822 | 0.0108 |
| Phoebis | **-0.1458** | -0.3630 | 0.0654 | 0.8126 | -0.0368 | -0.0915 | 0.0165 |
| Milanion | **-0.1488** | -0.3407 | 0.0387 | 0.8459 | -0.0375 | -0.0859 | 0.0097 |
| Antichloris | **-0.1490** | -0.3827 | 0.0762 | 0.8027 | -0.0376 | -0.0965 | 0.0192 |
| Cyclomia | **-0.1530** | -0.3144 | 0.0062 | **0.8910** | -0.0386 | -0.0793 | 0.0016 |
| Oraesia | **-0.1553** | -0.3526 | 0.0368 | **0.8505** | -0.0392 | -0.0889 | 0.0093 |
| Astraptes | **-0.1567** | -0.3104 | -0.0055 | **0.9080** | -0.0395 | -0.0783 | -0.0014 |
| Adelpha | **-0.1656** | -0.3466 | 0.0110 | **0.8852** | -0.0418 | -0.0874 | 0.0028 |
| Tigridia | **-0.1667** | -0.3600 | 0.0210 | **0.8725** | -0.0420 | -0.0908 | 0.0053 |
| Caligo | **-0.1693** | -0.3602 | 0.0161 | **0.8791** | -0.0427 | -0.0908 | 0.0041 |

| | Standardized Coefficients | | | | Unstandardized Coefficients | | |
|---|---|---|---|---|---|---|---|
| **Lepidoptera** | | | | | | | |
| **Genera** | **Esitmate** | **CI $l_{ower}$** | **CI$_{upper}$** | **Probability** | **Estimate** | **CI$_{Low}$** | **CI$_{High}$** |
| Myscelus | **-0.1695** | -0.3919 | 0.0437 | 0.8460 | -0.0427 | -0.0988 | 0.0110 |
| Desmia | **-0.1730** | -0.3511 | 0.0007 | **0.8992** | -0.0436 | -0.0885 | 0.0002 |
| Tarchon | **-0.1738** | -0.3430 | -0.0078 | **0.9102** | -0.0438 | -0.0865 | -0.0020 |
| Pachylia | **-0.1806** | -0.3751 | 0.0073 | **0.8909** | -0.0455 | -0.0946 | 0.0018 |
| Agaraea | **-0.1881** | -0.3765 | -0.0056 | **0.9067** | -0.0474 | -0.0949 | -0.0014 |
| Papilio | **-0.1915** | -0.3888 | -0.0016 | **0.9018** | -0.0483 | -0.0980 | -0.0004 |
| Dysschema | **-0.1961** | -0.3827 | -0.0159 | **0.9185** | -0.0494 | -0.0965 | -0.0040 |
| Apatelodes | **-0.2129** | -0.3964 | -0.0364 | **0.9391** | -0.0537 | -0.0999 | -0.0092 |
| Gonodonta | **-0.2186** | -0.3808 | -0.0606 | **0.9622** | -0.0551 | -0.0960 | -0.0153 |
| Hylesia | **-0.2266** | -0.4096 | -0.0512 | **0.9513** | -0.0571 | -0.1033 | -0.0129 |
| Pantographa | **-0.2532** | -0.4322 | -0.0816 | **0.9713** | -0.0638 | -0.1090 | -0.0206 |
| Dysodia | **-0.2554** | -0.4555 | -0.0672 | **0.9595** | -0.0644 | -0.1149 | -0.0169 |
| Emesis | **-0.2768** | -0.4933 | -0.0768 | **0.9630** | -0.0698 | -0.1244 | -0.0194 |

| | Standardized Coefficients | | | | Unstandardized Coefficients | | |
|---|---|---|---|---|---|---|---|
| **Lepidoptera** | | | | | | | |
| **Genera** | **Esitmate** | **CI $l_{ower}$** | **$CI_{upper}$** | **Probability** | **Estimate** | **$CI_{Low}$** | **$CI_{High}$** |
| Xylophanes | **-0.3059** | -0.4798 | -0.1401 | **0.9916** | -0.0771 | -0.1210 | -0.0353 |

**Table S2.**

| Network Property | 1997-2001 | 2012-2018 |
|---|---|---|
| **Node Richness** | | |
| Host plant | 109 [325] | 79 [216] |
| Herbivore | 32[941] | 24 [257] |
| Parasitoid | 10 [385] | 2 [67] |
| Total | 151 [1651] | 105 [540] |
| **Link Richness** | | |
| Host Plant-Herbivore | 442 [1654] | 199 [409] |
| Herbivore-Parasitoid | 83 [547] | 19 [80] |
| Total | 525 [ 2201] | 218 [489] |

**Table S3.**

| | Plant-Herbivore | Herbivore-Parasitoid |
|---|---|---|
| $\beta_{wn}$ | 0.92 | 0.81 |
| $\beta_{ST}$ | 0.27 | 0.68 |
| $\beta_{OS}$ | 0.64 | 0.14 |
| $\beta_{S}$ | 0.70 | 0.78 |

[1] $\beta$ calculated as Sorensen's Index

**Table S4.**

| | Climate Variable | Estimate | Std.Error | P.Value | R.squared |
|---|---|---|---|---|---|
| Annual Mean | Precip (mm) | 0.036 | 0.035 | 0.315 | 0.029 |
| | Tmin | 0.028 | 0.005 | 0.000 | 0.475 |
| | Tmax | 0.021 | 0.007 | 0.005 | 0.200 |
| | Avg. Temp. | 0.013 | 0.005 | 0.014 | 0.217 |
| Anomaly | Extreme Precip. Event | 0.011 | 0.034 | 0.749 | 0.003 |
| | Drought Events | -1.165 | 0.229 | 0.000 | 0.426 |
| | Tmin | 0.061 | 0.023 | 0.012 | 0.166 |
| | Tmax | 0.123 | 0.040 | 0.004 | 0.211 |
| | Avg. Temp. | 0.128 | 0.106 | 0.241 | 0.054 |
| Coefficient of Variation | Precip (mm) | 1.000 | 0.000 | 0.000 | 1.000 |
| | Tmin | 0.000 | 0.002 | 0.950 | 0.000 |
| | Tmax | 0.000 | 0.000 | 0.508 | 0.013 |
| | Avg. Temp. | 0.001 | 0.000 | 0.000 | 0.559 |

**Table S5**.

| | Climate Variable | Estimate | Std. Error | P.value | R.squared |
|---|---|---|---|---|---|
| **Wet Season** | | | | | |
| Annual Mean | Precip (mm) | 0.030 | 0.039 | 0.440 | 0.017 |
| | Tmin | 0.028 | 0.005 | 0.000 | 0.452 |
| | Tmax | 0.019 | 0.007 | 0.009 | 0.177 |
| | Avg. Temp | 0.013 | 0.005 | 0.011 | 0.230 |
| Anomaly | Days of Extreme Precip. | -0.002 | 0.032 | 0.942 | 0.000 |
| | Dry Days | -0.894 | 0.197 | 0.000 | 0.370 |
| | Tmax | -0.006 | 0.012 | 0.605 | 0.008 |
| | Tmin | 0.012 | 0.021 | 0.560 | 0.010 |
| | Avg. Temp | -0.032 | 0.013 | 0.023 | 0.190 |
| Coefficient of Variation | Precip (mm) | 1.000 | 0.000 | 0.000 | 1.000 |
| | Tmin | 0.001 | 0.002 | 0.760 | 0.003 |
| | Tmax | 0.000 | 0.000 | 0.220 | 0.043 |
| | Avg. Temp | 0.001 | 0.000 | 0.000 | 0.579 |
| **Dry Season** | | | | | |
| Annual Mean | Precip (mm) | 0.081 | 0.049 | 0.102 | 0.077 |

| | | | | | |
|---|---|---|---|---|---|
| | Tmin | 0.028 | 0.008 | 0.002 | 0.250 |
| | Tmax | 0.039 | 0.012 | 0.002 | 0.248 |
| | Avg. Temp | 0.014 | 0.010 | 0.165 | 0.079 |
| Anomaly | Days of Extreme Precip. | -0.001 | 0.009 | 0.881 | 0.001 |
| | Dry Days | -0.234 | 0.072 | 0.003 | 0.235 |
| | Tmax | -0.003 | 0.004 | 0.377 | 0.023 |
| | Tmin | 0.007 | 0.008 | 0.385 | 0.022 |
| | Avg. Temp | -0.005 | 0.005 | 0.331 | 0.039 |
| Coefficient of Variation | Precip (mm) | 1.000 | 0.000 | 0.000 | 1.000 |
| | Tmin | -0.015 | 0.008 | 0.059 | 0.101 |
| | Tmax | 0.001 | 0.000 | 0.007 | 0.201 |
| | Avg. Temp | 0.000 | 0.000 | 0.702 | 0.006 |

**Table S6.**

| Year | Plant | | Caterpillar | | Parasitoid | | Total Sampled | | Sample Effect/Hectare | Number of Volunteers per year |
|---|---|---|---|---|---|---|---|---|---|---|
| | Family | Species | Family | Species | Family | Species | Caterpillar | Parasitoid | | |
| 1997 | 37 | 73 | 19 | 111 | 6 | 51 | 726 | 64 | 234 | 22 |
| 1998 | 58 | 152 | 27 | 261 | 8 | 114 | 1859 | 237 | 350 | 51 |
| 1999 | 56 | 159 | 26 | 333 | 8 | 143 | 2531 | 273 | 270 | 31 |
| 2000 | 59 | 200 | 28 | 406 | 7 | 115 | 2673 | 136 | 346 | 50 |
| 2001 | 59 | 136 | 24 | 255 | 5 | 41 | 1603 | 48 | 306 | 40 |
| 2002 | 53 | 149 | 24 | 234 | 6 | 70 | 2422 | 105 | 306 | 40 |
| 2003 | 46 | 118 | 26 | 176 | 5 | 42 | 1820 | 51 | 318 | 43 |
| 2004 | 66 | 190 | 24 | 198 | 6 | 78 | 2595 | 91 | 266 | 30 |
| 2005 | 55 | 143 | 23 | 172 | 4 | 43 | 1662 | 84 | 266 | 30 |
| 2006 | 51 | 121 | 17 | 82 | 4 | 22 | 699 | 41 | 146 | 1 |
| 2007 | 61 | 163 | 21 | 163 | 6 | 39 | 1416 | 66 | 322 | 44 |

| | | | | | | | | | | |
|---|---|---|---|---|---|---|---|---|---|---|
| 2008 | 55 | 169 | 23 | 216 | 4 | 49 | 1747 | 65 | 186 | 10 |
| 2009 | 42 | 94 | 23 | 126 | 5 | 23 | 647 | 43 | 146 | 10 |
| 2010 | 28 | 45 | 16 | 64 | 3 | 5 | 289 | 3 | 146 | 1 |
| 2011 | 21 | 36 | 16 | 43 | 3 | 4 | 146 | 4 | 182 | 9 |
| 2012 | 8 | 18 | 11 | 17 | 2 | 3 | 152 | 2 | 182 | 9 |
| 2013 | 8 | 20 | 12 | 21 | 3 | 4 | 73 | 4 | 146 | 9 |
| 2014 | 14 | 18 | 12 | 20 | 3 | 3 | 31 | 2 | 174 | 7 |
| 2015 | 7 | 26 | 12 | 20 | 4 | 6 | 120 | 4 | 238 | 23 |
| 2016 | 10 | 13 | 8 | 11 | 3 | 3 | 24 | 3 | 198 | 13 |
| 2017 | 19 | 39 | 17 | 40 | 2 | 7 | 117 | 1 | 198 | 13 |
| 2018 | 48 | 147 | 24 | 153 | 3 | 41 | 685 | 30 | 146 | 13 |
